# Engineering paralog-specific PSD-95 synthetic binders as potent and minimally invasive imaging probes

**DOI:** 10.1101/2021.04.07.438431

**Authors:** Charlotte Rimbault, Christelle Breillat, Benjamin Compans, Estelle Toulmé, Filipe Nunes Vicente, Monica Fernandez-Monreal, Patrice Mascalchi, Camille Genuer, Virginia Puente-Muñoz, Isabel Gauthereau, Eric Hosy, Gregory Giannone, Ingrid Chamma, Cameron D. Mackereth, Christel Poujol, Daniel Choquet, Matthieu Sainlos

## Abstract

Despite the constant advances in fluorescence imaging techniques, monitoring endogenous proteins still constitutes a major challenge in particular when considering dynamics studies or super-resolution imaging. We have recently evolved specific protein-based binders for PSD-95, the main postsynaptic scaffold proteins at excitatory synapses. Since the synthetic binders recognize epitopes not directly involved in the target protein activity, we consider them here as tools to develop endogenous PSD-95 imaging probes. After confirming their lack of impact on PSD-95 function, we validated their use as intrabody fluorescent probes. We further engineered the probes and demonstrated their usefulness in different super-resolution imaging modalities (STED, PALM and DNA-PAINT) in both live and fixed neurons. Finally, we exploited the binders to enrich at the synapse genetically encoded calcium reporters. Overall, we demonstrate that these evolved binders constitute a robust and efficient platform to selectively target and monitor endogenous PSD-95 using various fluorescence imaging techniques.

## Introduction

Fluorescence microscopy constitutes nowadays an essential method for cell biologists to monitor the localization and function of most proteins. The discovery of the green fluorescent protein (GFP) and its application as a gene-fused reporter together with the progress that followed with the isolation and evolution of variants that span the close-UV to near-IR spectrum with various photo-physical and -chemical properties have largely contributed to the wide spreading of this approach (**Rodriguez *et al.*, 2017**). Alternative labelling methods such as those relying on engineered enzymes have further expanded the possibilities of imaging approaches by allowing the direct coupling of high-performance organic dyes (**Lavis, 2017**; **Xue *et al.*, 2015**). In parallel, technical breakthroughs in imaging methods have allowed to overcome the diffraction limit and are now enabling optical imaging of biological samples at the nanoscale (**Liu *et al.*, 2015**; **Sahl *et al.*, 2017**; **Schermelleh *et al.*, 2019**). However, while these advances have expanded the scope of application of fluorescence imaging techniques, they have also generated a pressing need for improved labelling strategies (**Choquet *et al.*, 2021**).

Indeed, the capacity to accurately investigate by fluorescence imaging the dynamics of endogenous proteins still constitutes a technical challenge. Antibodies, when available, can only be used against ectodomain-presenting proteins and still suffer from their large size and divalency. In parallel, the main drawbacks of most alternative labelling strategies for proteins (fluorescent protein, enzyme or tag genetic fusions) are associated with non-physiological regulation of the modified gene expression level and the potential impact of the fusion on the protein of interest function. Recent developments in gene editing methods(**Bukhari and Müller, 2019**) provide efficient means to circumvent the issue of expression level by directly modifying the endogenous gene, but their implementation is still not straightforward and furthermore intrinsically involves modifying the target protein with a fluorescent tag that can alter its function.

In this context, with the recent progress in directed evolution techniques, recombinant small domain binders capable of specifically recognizing endogenous proteins without impairing their function constitute a promising avenue for the development of minimally invasive labelling probes (**Bedford *et al.*, 2017**; **Helma *et al.*, 2015**). The increasing diversity in terms of validated molecular scaffolds, such as antibody fragments (scFv or V_H_H)(**Muyldermans, 2021**) or other domains (DARPins (**Boersma and Plückthun, 2011**), monobodies (**Sha *et al.*, 2017**), affimers (**Tiede *et al.*, 2017**), …), provides a large variety of randomized surfaces that can recognize and bind virtually any protein of interest. In addition to their recombinant nature, which facilitates their characterization and allows further engineering -notably to convert them into fluorescent probes- these tools importantly alleviate the need to directly alter the gene of interest. Additionally, their small size allows to bring fluorophores coupled to the engineered evolved domain in close proximity of the targeted protein for advanced imaging techniques.

Two recent studies (**Fukata *et al.*, 2013**; **Gross *et al.*, 2013**) have applied such a strategy to PSD-95, the major postsynaptic scaffold protein at excitatory synapses (**Chen *et al.*, 2005**; **Cheng *et al.*, 2006**), by evolving synthetic binders as key recognition modules for developing imaging probes. PSD-95 plays a key role in organizing receptors, ion-channels, adhesion proteins, enzymes and cytoskeletal proteins at excitatory synapses (**Won *et al.*, 2017**; **Zhu *et al.*, 2016**). As a consequence, up or down-regulation of PSD-95 results in critical alterations in synapse morphology and function (**Won *et al.*, 2017**). In particular, overexpression of fluorescent protein-fused PSD-95 for imaging purposes is phenotypically marked and leads to an increase in dendritic spine number and size, as well as frequency and amplitude of miniature excitatory postsynaptic currents (EPSC) and affects synaptic plasticity (**El-Husseini *et al.*, 2000**; **Nikonenko *et al.*, 2008**; **Zhang and Lisman, 2012**). PSD-95 constitutes therefore an ideal candidate for developing labelling strategies that do not affect the protein expression levels. By exploiting evolved binders, a single chain variable fragment (PF11) (**Fukata *et al.*, 2013**) and a ^10^FN3-derived domain/monobody (PSD95.FingR) (**Gross *et al.*, 2013**), the two groups have been able to directly label endogenous PSD-95. However, in both cases, the precise epitopes remain non-characterized and furthermore, one of the binders, PSD95.FingR, can also recognize SAP97 and SAP102, two closely related proteins (**Gross *et al.*, 2013**; **Li *et al.*, 2018**). The latter point may constitute a clear limitation and additionally, the lack of defined epitopes questions the possibility of PSD-95 function perturbation.

Using a phage display selection approach with a ^10^FN3-derived library, we have recently isolated and characterized three monobodies targeting PSD-95 (**Rimbault *et al.*, 2019**). The clones were targeted against PSD-95 tandem PDZ domains and showed remarkable specificity for PSD-95, in particular when considering the high sequence conservation of paralogs (SAP97, SAP102 and PSD-93). Importantly, all the clones recognized epitopes situated outside of the PDZ domains binding groove in regions not subjected to post-translational modifications. These properties represent a prerequisite to identify binders having with a minimal impact on the tandem domain function and consequently on the full-length protein. As such they constitute ideal candidates to engineer and develop minimally invasive imaging probes to monitor endogenous PSD-95.

We describe here the exploitation of specific PSD-95 binders as a platform to develop a series of labeling tools for the endogenous synaptic scaffold protein as well as excitatory synapses targeting modules. We first evaluated the potential impact of each evolved ^10^FN3 domain binding on PSD-95 function as well as their capacity to be exploited as intrabody-type imaging tools. The selected binders were further engineered to allow their use in various super-resolution imaging modalities (STED, PALM and DNAPAINT). Finally, beyond their direct exploitation as PSD-95 reporters, we validated the strategy to use their binding properties to enrich and address protein-based sensors to the postsynaptic density with the genetically encoded calcium reporter GCaMP6/7f (**Chen *et al.*, 2013**; **Dana *et al.*, 2019**). We termed the approach ReMoRA (Recombinant binding Modules for minimally invasive Recognition and Addressing of endogenous protein targets) as a sub-class of the intrabody general use with applications in fluorescence imaging where emphasis is set on the absence of interference with the targeted protein function.

## Results

### Impact of Xph15/18/20 on PSD-95 PDZ domains function

We have recently selected and isolated ^10^FN3-derived clones that bind to the tandem PDZ domains of PSD-95 (**Rimbault *et al.*, 2019**) (**Fig. 1a**). Three of the evolved ^10^FN3 domains, which displayed specific recognition of the target, were characterized in depth in particular with respect to the identification of their respective epitope. Two monobodies (Xph15 and Xph20) shared a similar epitope situated on PDZ domain 1 on the opposite side of the surface as compared to the canonical functional region of the domain. Indeed, as protein-protein interaction modules, the principal function of PDZ domains is to bind the C-terminus of their protein partner via a defined solvent exposed groove. The third monobody (Xph18) presented an extended epitope that spread on both domain 1 and 2, also in regions distant from the two binding grooves. As the three evolved binders did not directly block the two PDZ domains binding sites, we envisaged their use as minimally invasive targeting modules.

**Fig. 1.**
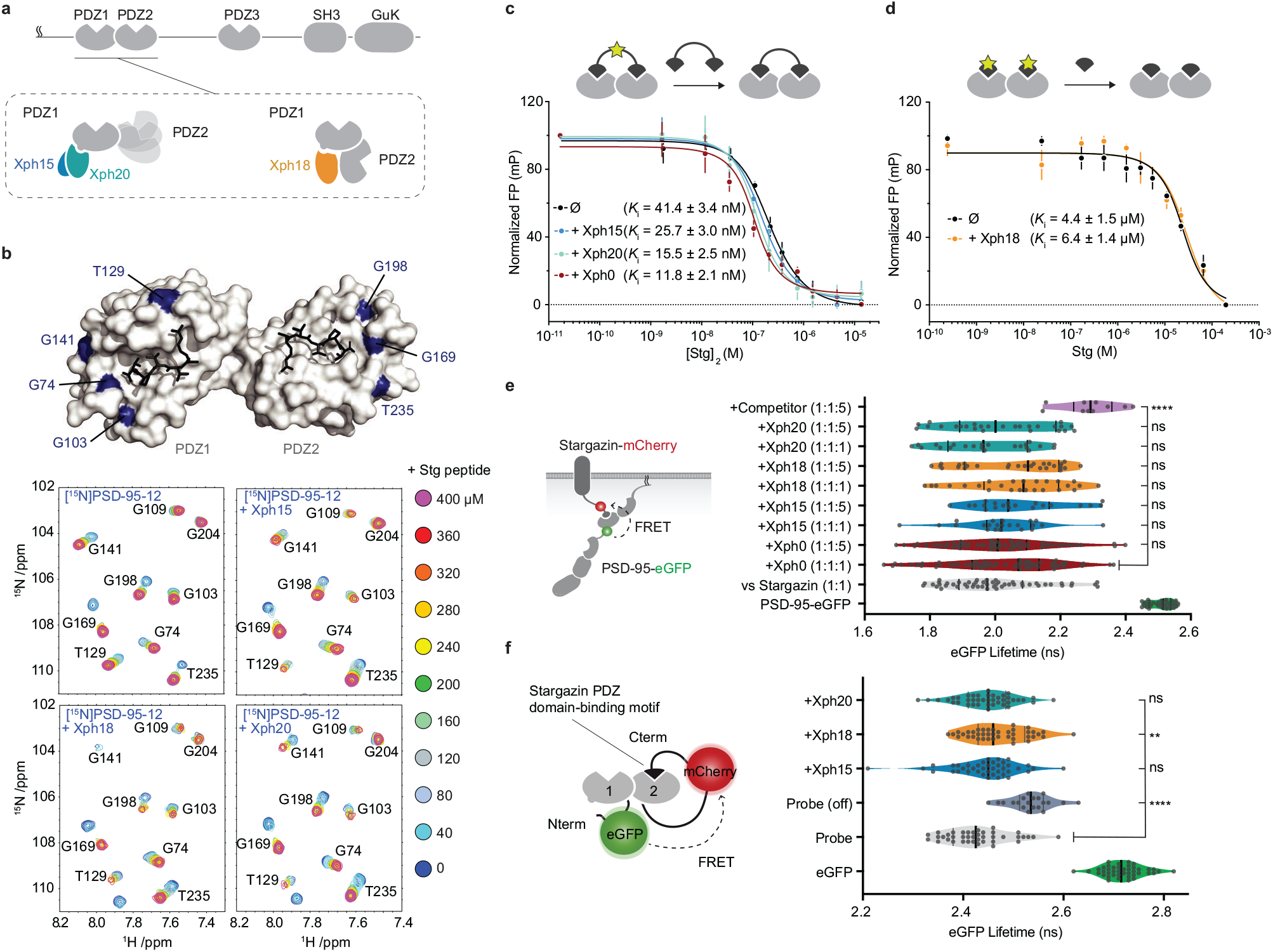
Evaluation of the impact of evolved ^10^FN3 domains binding on the PDZ domains function. **a**, PSD-95 domain organization and binding models of the three clones investigated. **b**, Titrations of a monovalent stargazin-derived peptide against PSD-95-12 in absence or presence of Xph15, Xph18 and Xph20. Surface representations of PSD-95 tandem PDZ domains (PDB ID 3GSL, domain 1 on the left and domain 2 on the right) with ligand modelled in (RTTPV derived from stargazin C-terminus and aligned from PDB ID 3JXT, black sticks) and with location of the residues annotated in the NMR titration spectra in blue: Gly74, Gly103, Thr129 and Gly141 report on stargazin binding to PDZ1; Gly 169, Gly 198 and Thr235 report on stargazin binding to PDZ2. Selected region of an overlay of ^1^H,^15^N-HSQC spectra corresponding to 200 μM of [^15^N]PSD-95-12 titrated with 0, 40, 80, 120, 160, 200, 240, 280, 320, 360, and 400 μM peptide ligand based on the C-terminus of stargazin (Stg) in the absence of evolved binder or in complex with 240 μM of Xph15, Xph18, or Xph20. Complete spectra are found in Supplementary Fig. 1. **c**, Competitive fluorescence polarization titrations between divalent stargazin-derived ligands and PSD-95-12 with or without Xph clones (5 μM each, mean ± SD of 3 independent titrations). **d**, Competitive fluorescence polarization titrations between monovalent stargazin-derived ligands and PSD-95-12 with or without Xph18 (20 μM, mean ± SD of 3 independent titrations). **e**, Lifetime of eGFP inserted in PSD-95 in presence of stargazin (acceptor-containing protein) and indicated constructs (molar ratio of DNA constructs specified as donor:acceptor:ligand). Violin plots show median, first and third quartile and all individual data points (each corresponding to a single cell) pooled from at least two independent experiments. Statistical significance determined by one-way ANOVA followed by Dunnett’s multiple comparison test. **f**, Lifetime of eGFP in a PSD-95-12-derived FRET reporter system in presence of indicated constructs (used at 5 molar equivalents of DNA compared to the FRET probe). Violin plots show median, first and third quartile and all individual data points (each corresponding to a single cell) pooled from at least two independent experiments. Statistical significance determined by one-way ANOVA followed by Dunnett’s multiple comparison test.

In an initial step prior to designing tools that target endogenous PSD-95, we sought to further characterize the binding properties of the three monobodies in the context of the tandem PDZ domains function. We first used in-solution NMR to evaluate if the PDZ domains binding properties to cognate ligands were affected by the presence of either of the clones (**Fig. 1b**, **Supplementary Fig. 1**). A peptide derived from the C-terminus of a known PSD-95 PDZ domain binder, the auxiliary AMPA receptor (AMPAR) subunit stargazin (Stg), was titrated against a ^15^N-labelled PSD-95 tandem construct containing PDZ domains 1 and 2. In addition, the peptide was titrated against the same ^l5^N-labelled PSD-95 construct pre-bound with either binding is generally unaffected by the presence of the Xph15, Xph18 or Xph20. A series of 2D ^l5^N-HSQC spectra binders. Conversely, by looking at residues at the PSD-95 were used to follow PSD-95 residues during each titration, and binder interface, the added Stg peptides also did not and in all cases residues on both PDZ1 and PDZ2 were able detectably affect the binding of the Xph monobodies to fully interact with the Stg peptide. Qualitatively, each of the reporter residues shown in **Fig. 1b** have similar titration behavior and final crosspeak positions in the ^l5^N-HSQC spectra, and therefore support the fact that the peptide binding is generally unaffected by the presence of the binders. Conversely, by looking at residues at the PSD-95 and binder interface, the added Stg peptides also did not detectably affect the binding of the Xph monobodies (Supplementary Fig. 1). These results confirm the simultaneous binding of both PDZ domain ligand and monobody and indicate that the PDZ domain binding properties are not detectably modified in the presence of the evolved Xph binders. In parallel, we also set up fluorescence polarization assays to determine the binding affinity of representative PSD-95 PDZ domain ligands in presence or absence of the monobodies. To this end, we used FITC-labelled peptides derived from the last 15 amino acids of stargazin as probes and the recombinant tandem PDZ domains 1 and 2. In order to minimize the effect of varying concentrations of the PDZ domains and the clones, we performed competition assays at constant concentrations of the monobodies, the PDZ domains, and the reporter probe. The potential effect of the evolved ^10^FN3 domain binding was first assessed using a divalent ligand titrated with a non-fluorescent divalent competitor both derived from stargazin as a model for complex multivalent interactions (**Fig. 1c and Supplementary Fig 2**) (**Sainlos et al., 2011**). Competitions performed in the absence of ligand or with a naïve clone (Xph0 that does not bind PSD-95) were similar to the ones obtained with Xph15 and 20. In contrast, the presence of Xph18 impaired binding of the fluorescent divalent probe. This effect was abolished, in agreement with the NMR observations, by the use of monovalent stargazin-derived probe and competitor (**Fig. 1d and Supplementary Fig 2**). These results suggest that the observed inhibitory effect results from the conformational constraints imposed on the two domains orientation by Xph18 binding rather than from the blocking or direct impairment of the PDZ domains binding ability.

**Fig. 2.**
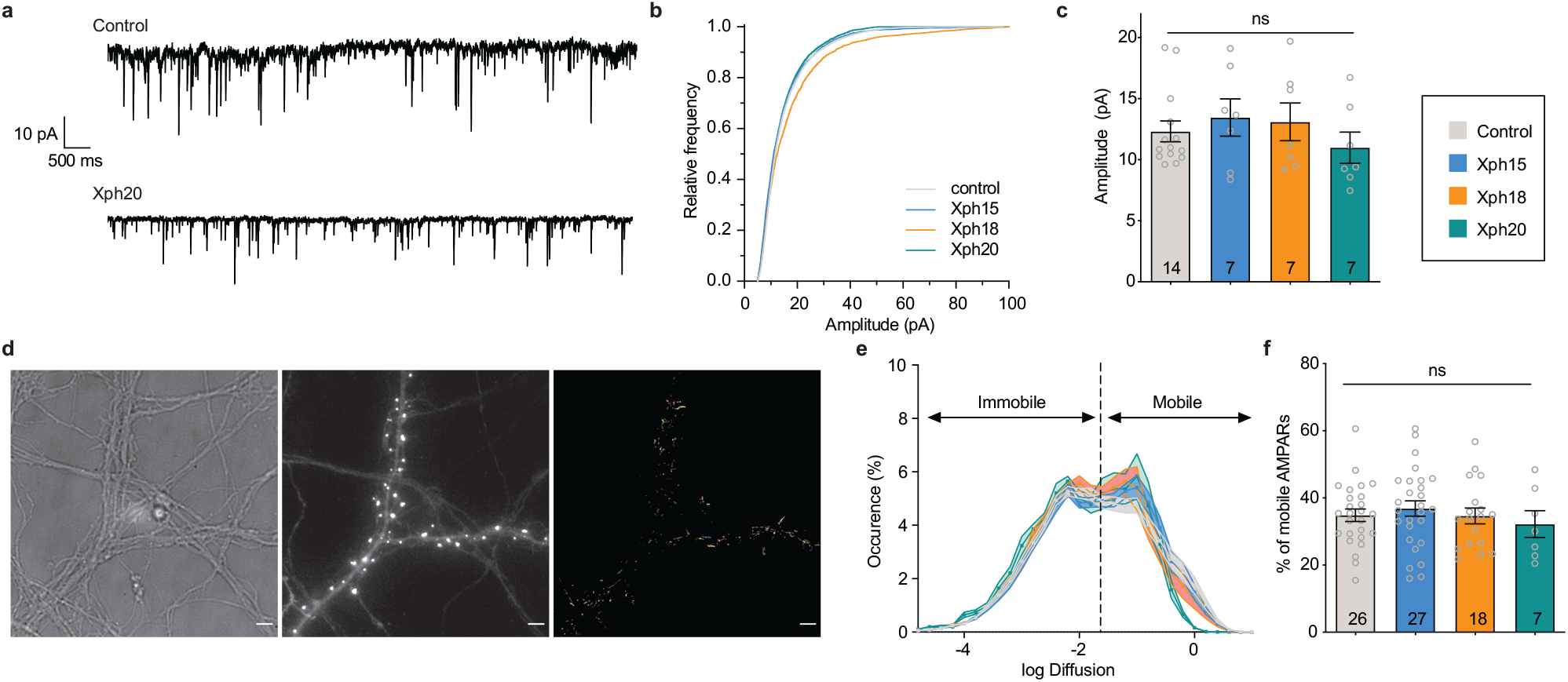
Evaluation of monobodies binding to endogenous PSD-95. **a**, Representative mEPSCs traces from pyramidal neuron primary culture, **b**, Cumulative distribution of miniature events amplitude, **c**, Average amplitude of mEPSCs (mean ± SEM, each dot represents a unique cell), d, Representative images for AMPARs single-particle tracking by uPAINT (left, DIC of neuron culture; middle, epifluorescence image of Xph20-eGFP expression pattern; right, trajectories; scale bar5 μm). **e**, Average distribution of instantaneous diffusion coefficients obtained by uPAINT of synaptic AMPAR with typical bimodal distribution. Error bars indicate cell-to-cell variability, **f**, Percentage of mobile AMPARs (mean ± SEM, each dot represents the mean value of mobile AMPAR per cell). Statistical analysis was performed with an ordinary one-way ANOVA test.

Together, the NMR study and the fluorescence polarization assay indicate that the binding of, on the one hand, Xph clones and, on the other, PDZ domain ligands are independent events that are not detectably affected by long-range conformational modifications. However, we note that due to the constraints imposed by Xph18 binding on the conformational flexibility of the two PDZ domains, certain complex interactions may be impaired.

Next, we investigated if these properties were preserved in complex cellular environments. We therefore evaluated by a FRET/FLIM assay in cell lines the binding of PSD-95 to its partners (represented here by stargazin) via its PDZ domains in presence and absence of the monobodies. We used both the recombinant full-length proteins (**Fig. 1e**) as well as a reporter system that focused on interactions mediated by PDZ domain1 and 2 (**Fig 1f**). In both cases, even at high molar ratio, we could not detect any significant effect on the measured donor lifetime associated with the binding of either Xph15, Xph18 or Xph20. For the full-length PSD-95 system, the median lifetimes obtained in the presence of the three clones, even at a 5-fold molar ratio in the transfected plasmids, were within the variability observed in the presence of a naïve clone (Xph0, between 2.0 and 2.1 ns). On the contrary, co-transfection of a soluble PDZ domain (PSD-95 2^nd^ PDZ domain, termed here competitor) clearly increased the lifetime to 2.3 ns. Results with the FRET probe based on PDZ domains 1 and 2 were comparable, with an absence of significant modification of the probe lifetime in the presence of the monobodies in comparison to a mutant of the probe in which the PDZ domain-binding motif was deleted (Probe off). A moderate effect was observed by statistical analysis in the case of Xph18, which could be attributed to the constraint imposed by its binding to both PDZ domain 1 and 2. In agreement with the NMR and fluorescence polarization experiments, these results therefore indicate that the primary function of PSD-95 PDZ domains as protein-protein interaction modules is not detectably affected by the interaction with any of the three synthetic binders in a model cellular environment.

### Impact of Xph15/18/20 on PSD-95 function

The main function of PSD-95 is to organize transmembrane receptors such as glutamate receptors at the postsynaptic density, and link them to intracellular signaling molecules. Among these, the PSD-95 interaction with AMPARs through the TARP auxiliary subunits has been particularly well characterized. We and others have previously shown that impairment of the interactions by genetic (**Bats *et al.*, 2007**) or chemical means (**Sainlos *et al.*, 2011**) resulted in a reduction of AMPAR synaptic currents and increased lateral mobility.

In order to rule out any possible effect of the monobodies on endogenous PSD-95 properties, we evaluated in hippocampal neuron primary cultures if the presence and binding of the Xph monobodies could impact AMPAR organization and function. To this end, and anticipating exploitation of the evolved ^10^FN3 domains as fluorescence imaging tools, we expressed the clones as fusions to eGFP in association with the expression regulating system developed for the abovementioned PSD95.FingR (**Gross *et al.*, 2013**). The probe regulation is achieved by fusion of a transcription repressor and a zinc finger in combination with the incorporation of the corresponding zinc finger-binding motif upstream of the reporter gene in the expression plasmid (**Supplementary Fig 3**). In this system, while eGFP is used to monitor the binding module and its target, the regulation system allows to avoid overexpression of the synthetic binder compared to its endogenous target.

**Fig. 3.**
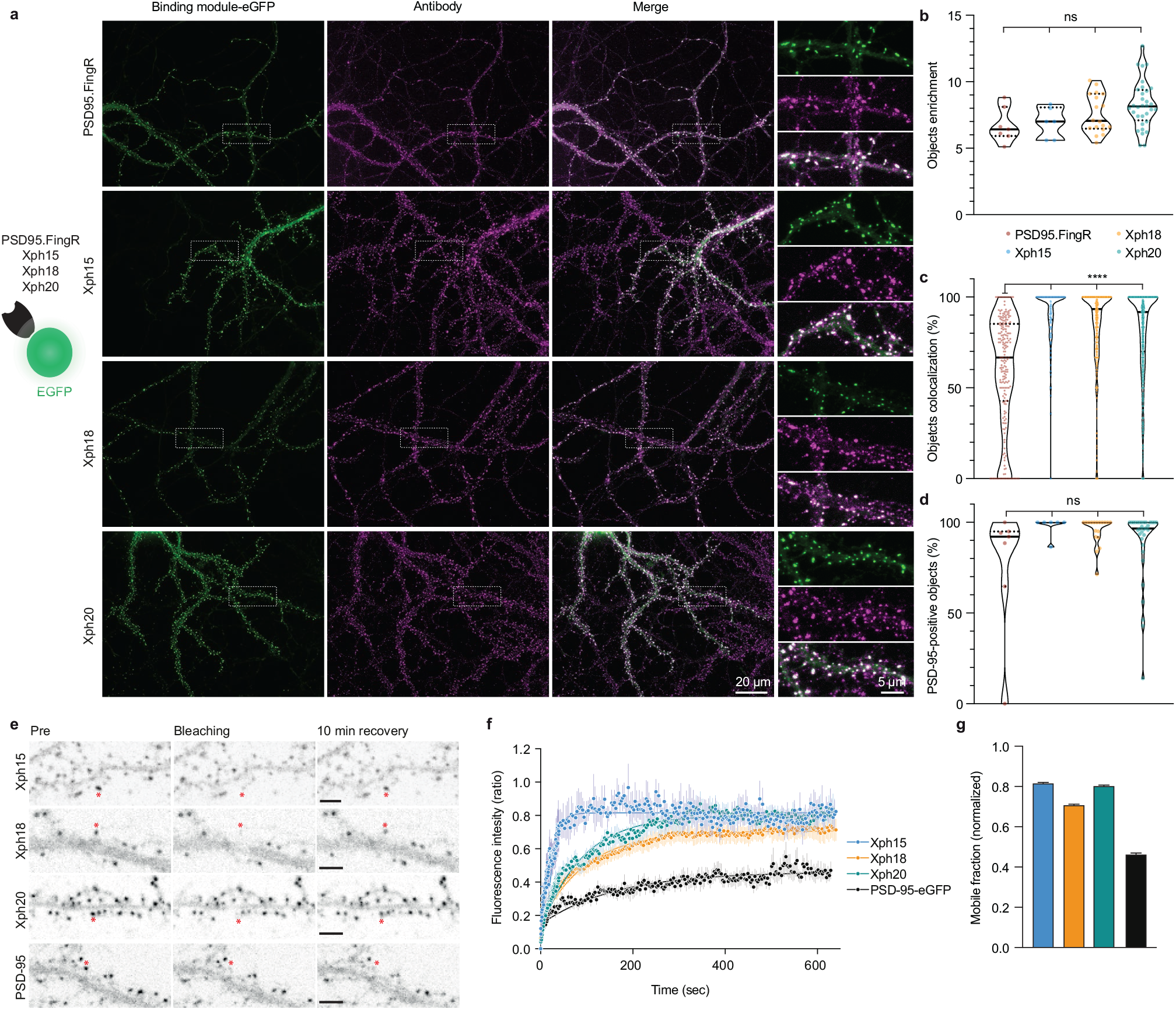
Evaluation of monobodies as fluorescent reporter probes, **a**, Representative epifluorescence images of the eGFP-fused binding modules vs immunostaining of endogenous PSD-95 domain. For the zoomed regions, top: binding module; middle: antibody staining; bottom: merge, **b**, Enrichment of object vs shaft fluorescence signal. Violin plots show median, first and third quartile and all individual data points (each corresponding to the analysis of a single acquired image) pooled from at least two independent experiments, **c**, Percentage of eGFP vs antibody objects colocalization (obtained by determining the % of common pixels within a probe labeled object with PSD-95 immunostaining). Violin plots show median, first and third quartile and all individual data points (each corresponding to a detected object) pooled from at least two independent experiments, **d**, Percentage of PSD-95 positive objects defined as objects with more than 50% pixel in common. Violin plots show median, first and third quartile and all individual data points (each corresponding to the analysis of a single image) pooled from at least two independent experiments, **e**, Representative images for FRAP experiments with eGFP fusion proteins, the red asterisk indicating the bleached dendritic spine. Scale bars 5 μm. **f**, Fluorescence recovery analysis (mean ± SEM with fitted curve, n= 6, 11, 9 and 7 spines for Xph 15, Xph18, Xph20 and PSD-95-eGFP respectively), **g**, Mobile probe fraction (mean ±SEM, n and color code same as 1).

We first investigated if the AMPAR-mediated synaptic currents were affected by the presence of the various monobodies. Comparison of control non-transfected neurons with neurons transfected with Xph15, Xph18, or Xph20 did not reveal any significant difference on spontaneous miniature excitatory postsynaptic currents (mEPSCs, **Fig. 2a, b and c**). The amplitude distributions were highly similar (**Fig. 2b**) for all the monobodies except for a slight shift observed for Xph18 that we attribute to cellular diversity. In line with these results, the mean amplitude values were not modified by the presence of any of the PSD-95 binders (**Fig. 2c**, control: 12.3 ± 3.2 pA (n = 14); Xph15: 13.4 ± 4.0 pA (n = 7); Xph18: 13.1 ± 4.1 pA (n = 7); Xph20: 11.0 ± 3.4 pA (n = 7); mean ± SEM with P>0.58 for the three clones using ordinary one-way ANOVA). We selected Xph15 and Xph18 as representative binders of the two types of epitopes (Xph15 and Xph20 share the same) and further investigated the miniature events. Similarly to the amplitude, nor the frequency, the rise time, orthe decay time were significantly modified as a result of the expression of the monobodies (**Supplementary Fig 4**).

**Fig. 4.**
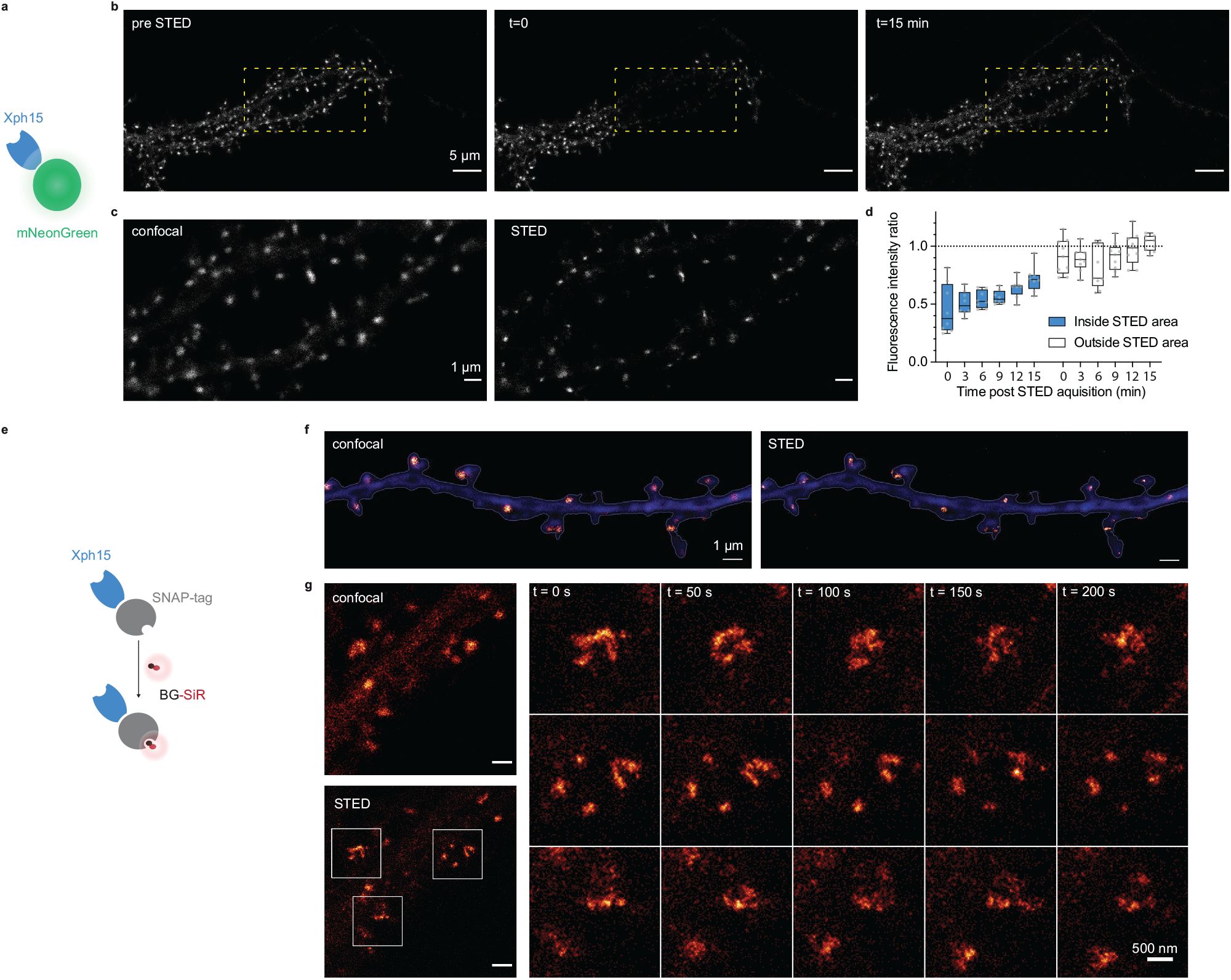
Evaluation of probes for STED imaging, **a**, Schematic representation of fluorescent protein-fused STED probe, **b**, Representative confocal images of a neuron transfected with Xph15-mNeonGreen before and after STED. The yellow box corresponds to the STED region, **c**, Confocal and STED images of the yellow box region from **b**. **d**, Evolution of fluorescence intensity over time of fluorescent objects subjected or not to STED (n = 8 and 9 for regions outside and inside of STED area, respectively). Box plots show median, first and third quartile, with whiskers extending to the minimum and maximum and all individual data points (each corresponding to a single object) pooled from at least two independent experiments, **e**, Schematic representation of SNAP-tag-fused STED probe with the BG-SiR fluorophore. **f**, Confocal and STED images of a neuron transfected repeated STED acquisitions with Xph15-SNAP-tag/BG-SiR. Time course of repeated STED acquisitions with Xph15-SNAP-tag/BG-SiR.

In addition to the electrophysiological measurements as an indicator of the proper synaptic recruitment of AMPARs, we also tested possible interference of the clones on the lateral mobility of surface AMPARs, as PSD-95 is the main AMPAR stabilizer (**Bats *et al.*, 2007**). Transfected and non-transfected culture neurons were sparsely labeled in live condition with an ATTO-647N-conjugated antibody against the GluA2 subunit ectodomain. Single-particle tracking was performed by using the uPAINT method (**Giannone *et al.*, 2010**) in order to gain insight on the AMPAR dynamics (**Fig. 2d**). In agreement with the absence of modification of excitatory currents, no detectable effect was observed for Xph-expressing neurons vs control non-transfected ones on the lateral mobility of surface AMPARs. The distributions of diffusion coefficients were highly similar for all conditions (**Fig. 2e**). Importantly, the percentage of mobile AMPARs was not increased in presence of any of the clones as could have been expected from a binder that would have perturbed interactions with either of the first two PDZ domains (**Fig. 2f**, control: 34.9 ± 9.5% (n = 26); Xph15: 36.9 ± 11.9% (n = 27); Xph18: 34.6 ± 9.9% (n = 18); Xph20: 32.2 ± 10.5% (n = 7); mean ± SD with P>0.73 by ordinary one-way ANOVA).

Altogether these experiments indicate that the binding of neither Xph15, Xph18, nor Xph20 affects endogenous PSD-95 function in its native environment as judged by the absence of impact on AMPAR properties. These results are therefore consistent with the nature of each clone’s respective epitope, which are found on regions of PSD-95 not involved in the PDZ domains binding of native cellular protein partners.

### Evaluation of Xph15/18/20 as endogenous PSD-95 imaging probes

The absence of any detectable effect of Xph clone binding on PSD-95 function constituted an obligatory first criterion to consider their use as a non-interfering imaging probe. As the three monobodies comply with this criterion (albeit with some reservation for Xph18), we next focused on confirming their capacity to label endogenous PSD-95 and on evaluating their specific properties as fluorescent probes.

First, we assessed the ability of Xph15,18 and 20 to bind and target a fluorescent protein to PSD-95 in primary hippocampal neuron culture. Neurons were transfected with the previously tested Xph-eGFP fusions (or PSD95.FingR-eGFP (**Gross et *al.*, 2013**), from which the expression vector was derived, as a comparison) chemically fixed after 23-27 days in vitro (DIV) and immunostained for PSD-95 (**Fig. 3a**). For all the binders tested, the eGFP signal was similarly strongly enriched on dendrites at postsynapse-like structures. The objects we observed presented in all cases a mean intensity enrichment ratio compared to the rest of the dendrite around 7 (**Fig. 3b**; PSD95.FingR: 6.7 ± 1.3; Xph15: 6.9 ± 1.1; Xph18: 7.6 ± 1.5; Xph20: 8.3 ± 1.8; mean ± SD with P>0.99 by ordinary one-way ANOVA followed by Tukey’s multiple comparison tests). This indicates that the four binders behave similarly in their capacity to address a fluorescent protein reporter to specific regions in neuronal cells. We next analyzed in each case how these objects colocalized with the labeling obtained by immunostaining of endogenous PSD-95 (**Fig. 3a, c and d**). In general, colocalization percentage values ranged from 0 to 100, which we attribute to the inherent differences of the two staining methods being compared *(i.e.* expressed reporter vs antibody labeling post-fixation and permeabilization). The colocalization of PSD-95 positive objects detected by antibody-immunostaining with the puncta revealed by the four investigated probes was strong (**Fig. 3d**, median > 90%, P > 0.14 by one-way ANOVA followed by Tukey’s multiple comparison tests), in agreement with reported values for PSD-95.FingR (**Cook *et al.*, 2019**; **Gross *et al.*, 2013**). However, the three Xph clones clearly showed a stronger enrichment in high colocalization values compared to PSD95.FingR (**Fig. 3c**, PSD95.FingR: 60.1 ± 30.7; Xph15: 91.5 ± 14.7; Xph18: 84.9 ± 21.7; Xph20: 81.4 ± 24.0; mean ± SD with P < 0.0001 for PSD95.FingR vs the other binders by ordinary one-way ANOVA followed by Dunnett’s multiple comparison test). We interpret this difference as a direct benefit from the specificity of the Xph clones for PSD-95 while PSD95.FingR, which can also bind SAP97 and SAP102, may also report to some extent the presence of these two paralog proteins. Considering the generally strong overlap of the GFP signal with the immunostaining of endogenous PSD-95, we conclude that the three monobodies label PSD-95 efficiently.

In order to evaluate the flexibility/versatility of the labeling system, we considered other fluorescent proteins, and in particular, a red fluorescent protein. We chose the recently described mScarlet-I as one of the brightest red reporters (**Bindels *et al.*, 2017**). Despite several attempts, we failed at expressing the Xph20-mScarlet-I fusion in transfected cultured neurons as a result of a toxicity not observed for the eGFP constructs. Transfer of the Xph20-mScarlet-I fusion into a non-regulated plasmid resulted in non-toxic expression of the probe, albeit at a higher level compared to PSD-95 endogenous expression levels. It therefore led to a homogenous filling of the whole neurons volume (**Supplementary Fig 5**). This indicates that the toxicity is here a consequence of the association of mScarlet-I with the regulation system. Replacement of mScarlet-I with another bright monomeric red fluorescent protein, mRuby2 (Lam *et al.*, 2012), abolished the observed toxic effect and provided a similar staining as compared to Xph-eGFP fusions (**Supplementary Fig 5**).

**Fig. 5.**
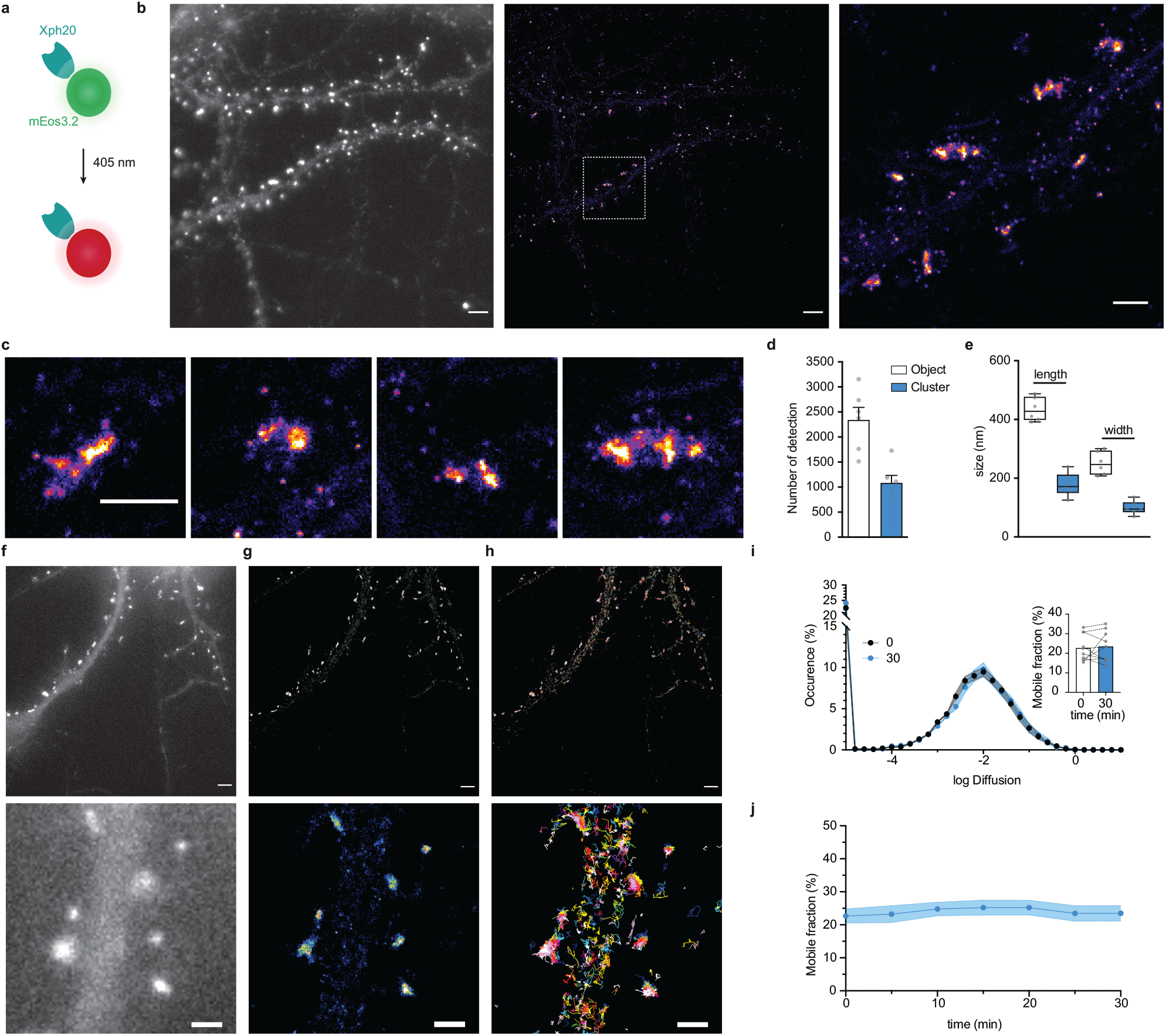
Evaluation of mEos3.2-derived probes for PALM and spt-PALM applications, **a**, Schematic representation of mEos3.2-fused probe, **b**, Representative epifluorescence and PALM images of Xph20-mEos3.2 in fixed culture neurons. Left: epifluorescence image obtained from the native non-photoconverted green form of mEos3.2; middle: super-resolution image obtained by PALM from a sequence of 20,000 images of sparse single molecules of the photoconverted red from of mEos3.2; right: zoomed region. Scale bar 5 μm. **c**, examples of individual synapses showing PSD-95 organization at the postsynaptic density (“object“) and sub-synaptic domain (“cluster“), d, Number of detections in “objects” vs “clusters” (mean ± SEM, each data point represents a single neuron), **e**, Morphological analysis of “objects” and “clusters” (mean ± SEM, each data point represents a single neuron). **f,g,h**, Representative epifluorescence and spt-PALM images of live culture neurons expressing Xph20-mEos3.2. epifluorescence of the expressed probe (before photoconversion) **(f)**, super-resolution intensity map obtained by sptPALM from a sequence of 4 000 images of sparse single molecules of the photoconverted red from of mEos3.2(**g**) and trajectories of PSD-95 tagged with Xph20-mEos3.2(**h**). Scale bars 5 and 2 μm for top and bottom images respectively, **i**, Average distribution of instantaneous diffusion coefficients obtained by spt-PALM of PSD-95 labeled with Xph20-mEos3.2 (at 0 min, to, beginning of the imaging session) or after 30 min of imaging (t;;o). Error bars indicate cell-to-cell variability. Insert: Percentage of the mobile fraction of probes at to vs t;;o (mean ± SEM, each dot represents a single cell, n = 10). **J**, Time course of the percentage of mobile probes (mean ± SEM, each dot represents a single cell, n = 10).

While these surprising results suggest that not all fluorescent proteins are compatible with the expression regulation system, they also highlight the critical need to match the target expression levels for imaging applications. In particular, as reported for PSD95.FingR, the expression regulation system applied to the Xph binders allows for long expression schemes without excessive or detectable over-production of the probe. This possibility in turn provides flexibility to handle the timing of the genetically encoded probe delivery without compromising the achieved labeling steady state.

The binding kinetics of the Xph clones previously evaluated by surface plasmon resonance (SPR) showed different but overall rather fast association and dissociation rate constants indicating fast exchanging complexes (half-lives of ∼2, 28, and 10 s for Xph15, Xph18 and Xph20 respectively for the isolated recombinant PSD-95 PDZ domains 1 and 2) (**Rimbault *et al.*, 2019**). These kinetic profiles were also associated with moderate affinities with binding constants in the low micromolar (4.3 and 2.6 μM for Xph15 and 18 respectively) to sub-micromolar range (330 nM for Xph20). We therefore sought to further evaluate how these properties would translate in the context of their use as PSD-95 labelling tools. To this end, we used fluorescence recovery after photobleaching (FRAP) to determine how the different probes interact with their target in its native environment. Fluorescence recovery was measured in photobleached single synapses (Xph objects) in neurons expressing the Xph-eGFP fusions (**Fig. 3e**). As expected, the 3 monobodies showed fast exchange rates ((**Supplementary Table 1**)) as well as high mobile fractions (**Fig. 3f** and (**Supplementary Table 1**); 81, 71 and 80% for Xph15, Xph18, and Xph20 respectively) compared to values reported in basal conditions for PSD-95-GFP knock-in (∼10% after 60 min (**Fortin *et al.*, 2014**)) or to the values obtained here with a transfected PSD-95-eGFP (46 %). The measured mobile fractions and the half-lives (∼10, 70, 60 s for Xph15, Xph18 and Xph20 respectively) are consistent with the SPR kinetics measurements with Xph15 being the fastest and Xph18 the slowest. We note that the results we obtained here for the probes account for the behavior of both the free and the PSD-95-bound populations. However, considering the large difference between the values obtained for the probes and for PSD-95, we can reasonably conclude that the Xph-derived probes exchange and are being renewed at a faster rate than their target.

Overall, this ensemble of results demonstrates first that Xph15, Xph18, and Xph20 can be used to efficiently recognize and bind endogenous PSD-95 with minimal impact on its function. In addition, the fusion of a fluorescent protein to the monobodies (together with the use of an expression regulation system) allows in this context to dynamically label PSD-95 in live conditions. Considering the large epitope recognized by Xph18, which as we have shown leads to a constrained conformation of the tandem PDZ domains, we chose here to only focus on Xph15 and Xph20 that both recognize a smaller epitope restricted to PDZ 1 to further develop the imaging tools and fully ensure minimal invasiveness of the resulting probes.

### Engineering probes for super-resolution imaging

The previous experiments validated the use of Xph-derived constructs as imaging probes to monitor endogenous PSD-95. The specific recognition properties of the evolved ^10^FN3 domains coupled to the capacity to match their expression levels to those of PSD-95 therefore provide an ideal platform to further elaborate our clones into more advanced probes, in particular for super-resolution imaging (SRI) applications. To this end, we modified the GFP reporter part of the probes with systems better adapted for various SRI modalities.

We first investigated how the probes performed with stimulated emission depletion (STED) microscopy. STED is a point-scanning method that relies on the simultaneous use of both an excitation and a depletion laser beam (**Sahl *et al.*, 2017**; **Vicidomini *et al.*, 2018**). Since the technique is compatible with a number of fluorescent proteins, its implementation is here relatively straightforward. A first attempt to determine if expression levels were compatible with STED imaging was performed on fixed cultured neurons expressing Xph20-eGFP. Without the need to improve the fluorescent protein part, the results were satisfactory with a clear gain in resolution when comparing STED and confocal imaging **(Supplementary Fig. 6a-b).**

**Fig. 6.**
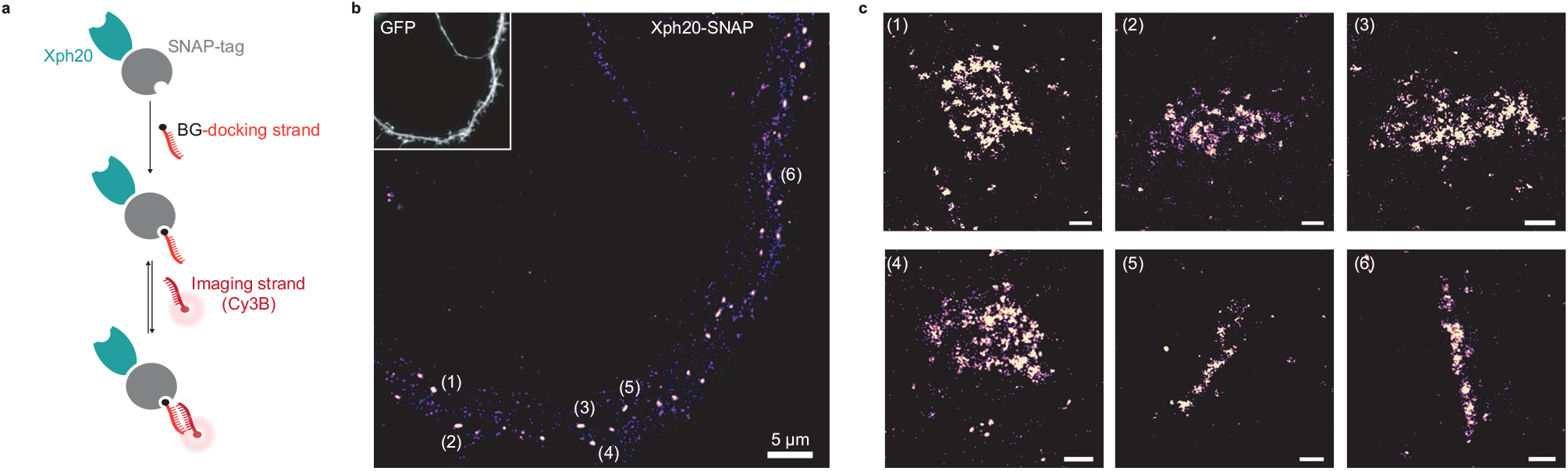
Evaluation of SNAP-tag-derived probes for DNA-PAINT super-resolution microscopy, **a**, Probe design and labeling scheme (BG: benzylguanine). **b**, Reconstructed DNA-PAINT image (10 Hz, 32000 frames) of Xph20-SNAP-tag in the dendrites of a DIV 14 hippocampal primary neuron (inset corresponding to soluble GFP). **c**, Magnified views of the regions marked in **b** (scale bars 100 nm).

In comparison to other imaging techniques, one of the main advantages of STED is the compatibility with live imaging, in particular for dynamic studies. While alternative methods exist to label endogenous PSD-95 post-fixation/permeabilization, live labeling of PSD-95 still remains a challenge for which Xph15 or Xph20 can provide solutions. A major drawback of STED is the high illumination intensities required in particular for efficient depletion that often results in photobleaching of the fluorophore. In this context, the fast exchanging properties of the probes could be an advantage and allow, by fast renewal of the probes, repeated acquisitions of the same region.

For live experiments, we therefore used the fastest exchanging binder, Xph15, to fully benefit from maximal probe replacement. In parallel, the fluorescent protein eGFP was replaced by mNeonGreen (**Shaner et al., 2013**), for its higher quantum yield and improved photostability. The Xph15-derived probe expressed well and provided a labeling similar to the eGFP construct in live dissociated neurons as judged by confocal microscopy (**Fig. 5a-c**). Despite the improved properties of mNeonGreen, application of a STED illumination invariably led to significant photobleaching of the area investigated (**Fig. 4b**). Nevertheless, the imaged area was repopulated over time with fresh probes as could be anticipated from the FRAP experiments. About 70% of the initial fluorescence was recovered in less than 15 minutes (**Fig. 4d**), allowing the area to be efficiently imaged repetitively. We note however that while the confocal imaging quality was comparable to the one obtained prior to the STED imaging, in that time scale, the STED quality was still noticeably degraded due to the loss of signal. Avoiding the repeated confocal imaging as well as reducing the area of STED imaging could be simple strategies to further improve the fluorescence recovery by limiting the photobleaching associated with unnecessary light exposition and by locally increasing the pool of intact probes vs photodamaged ones.

In order to more efficiently circumvent the loss of signal and enable faster repeated STED acquisitions, for instance with 3D-stacks or minute-timescale super-resolution investigations, we modified our strategy and opted for the use of brighter and more photoresistant organic dyes. To effectively functionalize our probes with such dyes, we replaced the fluorescent protein by a SNAP-tag (**Keppler *et al.*, 2003**) (**Fig. 4e**) and used a cell-permeant silicon rhodamine fluorophore (SiR) (**Lukinavičius *et al.*, 2013**) coupled to benzylguanine added prior to the imaging session. The SNAP-tag probe behaved comparably to the eGFP version, and synaptic objects, hallmark of PSD-95 neuronal distribution, could be visualized (**Fig. 4f**). Efficiency of the STED imaging was improved by the use of the brighter SiR dye (**Fig. 4f-g** and **Supplementary Fig. 6c-d**). Photostability and dynamic exchange of the probe allowed to perform timelapse acquisitions at about a 1-minute (50 s) frequency with minimal impact on the STED signal (**Fig. 4g**) thereby illustrating the advantage of organic dyes over fluorescent proteins for such applications.

Single-molecule localization microscopy (SMLM) is another strategy used to access spatial resolution below the limit imposed by the diffraction of light (**Sauer and Heilemann, 2017**). It relies on temporal decorrelation of fluorophore emissions to obtain sparsely located fluorescent entities while keeping the majority of the population in non-emissive states. One strategy to perform SMLM is photoactivation localization microscopy (PALM) based on the use of photoactivatable of photoconvertible fluorescent proteins. To implement this imaging modality, we thus replaced eGFP with the monomeric photoconvertible protein mEos3.2 (**Zhang *et al.*, 2012**). We considered Xph20 as the binding module for its stronger affinity and slower off-rate kinetics. The photoconvertible probe was expressed in dissociated cultured neurons and provided in its basal green state a labeling similar to the eGFP probe (**Fig. 5a and b**).

Stochastic photoconversion of mEos3.2 was first performed in fixed neurons, and analysis of the resulting image stacks used to generate super-resolved images (**Fig. 5b**). Efficiency of the fixation step on the probe was assessed by determination of the diffusion coefficients distribution of the detected single emitters. The results confirmed that a large majority of the investigated emitters were immobile (**Supplementary Fig. 7a-d**). The reconstructed maps showed a clear enrichment of the probe at synapses as already observed with diffraction limited imaging techniques and STED. Further analysis of the synaptic objects using the Tesselation-based segmentation method (**Levet *et al.*, 2015a**) revealed a non-homogenous distribution with the presence of higher density clusters. The clusters represented about half the number of detections measured for the whole synaptic objects. A tentative estimation of single emitters contribution (**Fig. 5d**, **Supplementary Fig. 7e**, median ∼10 detections) suggests the presence of ∼200-300 probe copies per synaptic objects. This value is consistent with reported estimations of PSD-95 synaptic copies (**Chen *et al.*, 2005**; **Sugiyama *et al.*, 2005**) and therefore suggests a close to stoichiometric labeling of the endogenous protein. Morphological analysis of the objects and clusters provided dimensions consistent with previous reports for PSD-95-mEos2 fusions with PALM (**Nair *et al.*, 2013**) or by STORM by labeling endogenous PSD-95 with antibodies (**Compans *et al.*, 2019**) (length: 434.5 ± 39.9 and 178.0 ± 39.3 nm; width: 251.1 ± 39.2 and 99.1 ± 22.4 nm for the objects and clusters respectively, **Fig. 5e**). Together, these results indicate that Xph20-mEos3.2 provides an accurate snapshot of PSD-95 nanoscale distribution in fixed samples.

**Fig. 7.**
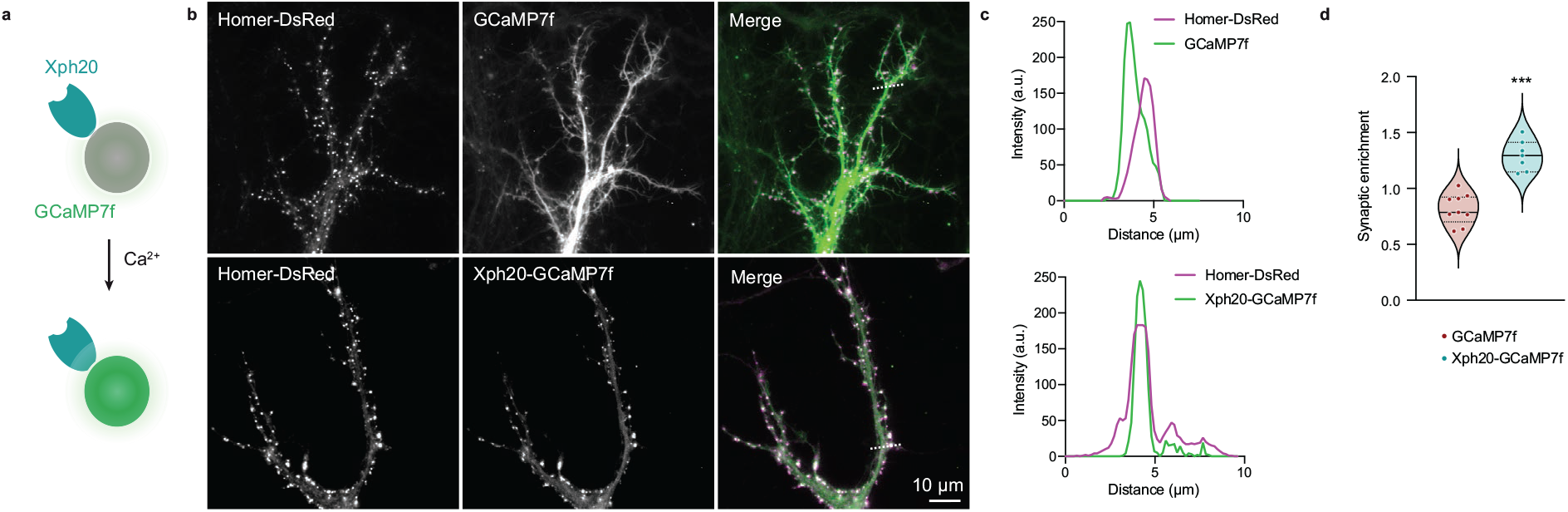
Application of the ReMoRa method for the synaptic targeting of calcium reporters, **a**, Schematic representation of calcium signaling probe, **b**, Comparison of the expression profile of targeted and regulated (Xph20-GCaMP7f, bottom panel) vs parental calcium sensor (GCaMP7f, top panel) for GCaMP7f synaptic targeting (GCaMP in green and Homer-DsRed in magenta in the merged images), **c**, Line scans from b comparing the probe repartition between shaft and spine compartments. The line scans show a clear enrichment of the regulated probe in neuronal spines, **d**, Probes synaptic enrichment determined using Homer-DsRed staining as a synaptic marker (n= 9 and 7 cells for GCaMP7f and Xph20-GCaMP7f respectively, from two independent experiments, P = 0.0002 by Mann-Whitney’s test).

PALM can also be performed on live samples, and single-particle tracking approaches yield in this case information on protein dynamics. This approach is typically achieved with a direct genetic fusion between the protein of interest and a photoconvertible fluorescent protein. Considering the efficiency of the evolved ^10^FN3 domain mediated labeling, we investigated here how this approach could be implemented with Xph20-mEos3.2. Indeed, single emitters are tracked on a time scale over an order of magnitude faster (∼500 ms) than the probe-target exchange dynamics (half-life of ∼10 s), which should avoid bias linked to the occurrence of particles alternating between PSD-95 bound and unbound states.

Tracked particles were detected within the whole dendrite (**Fig. 5f-h**). As observed previously with other imaging techniques, a strong enrichment of the probe was observed at the synapse when reconstructing the intensity maps corresponding to the accumulation of track coordinates. The probe diffusion coefficient showed a gaussian-like distribution, which suggests a homogenous population, with ∼80% of particles confined or immobile and only ∼20% mobile (**Fig. 5i**). These results are highly similar to those obtained with a mEos2-fused (**Chazeau *et al.*, 2014**) or a mVenus-fused PSD-95 (**Fortin *et al.*, 2014**) suggesting that we are essentially detecting probes bound to PSD-95. Indeed, a freely diffusive emitter, such as an unbound probe, would be characterized by faster diffusion coefficients (**Chazeau *et al.*, 2014**), that could not be detected in these experimental conditions. Importantly, single-particle tracking-PALM measurements could be repeated over the course of 30 minutes without detectable differences in the diffusion distribution (**Fig. 5j**) demonstrating that the method is robust and compatible with hour time scale live investigations such as for instance needed for synaptic plasticity events.

Considering the compatibility of our probes with STED and PALM, we next investigated their implementation to more recent super-resolution imaging techniques adapted to the detection of multiple distinct targets. DNA-PAINT (**Jungmann et al., 2014**) constitutes a powerful alternative approach to STORM or PALM for SMLM, in particular for multiplexing applications, as it allows sequential imaging of different proteins of interest with the same fluorophore. The technique is based on the use of pairs of short complementary oligonucleotides, one strand bound to a target or its probe (docking strand) and the other functionalized with a fluorophore (imager strand), that undergo fast dynamic exchange between the bound and unbound states. In order to couple the docking strand to the Xph-derived probes, we considered here the use of either SNAP- or HaloTag (**Los et al., 2008**) to enzymatically create a covalent bound with benzylguanine- or haloalkane-derived oligonucleotides (**Schlichthaerle et al., 2019**). For the binding module, as for PALM, we chose Xph20 for its stronger affinity.

Each construct was co-transfected with a soluble eGFP marker in dissociated culture hippocampal neurons and used to implement the DNA-PAINT method after chemical fixation. The self-labeling tags were each reacted with the corresponding docking strand and after removal of the excess material, the Cy3B-derived imaging strand was applied to the samples. A control experiment in which no docking strand was added confirmed the very low level of non-specific binding of the imaging strand in our conditions (**Supplementary Fig. 8a**). For the HaloTag fusion, a homogenous staining of the transfected neurons was observed (**Supplementary Fig. 8b**), suggesting a failure either of the target recognition or of the regulation system with this particular engineered enzyme. The reason for this failure was not investigated further. We note that the larger size of the HaloTag (34 kDa **vs** 20 or 27 kDa for the SNAP-tag and fluorescent proteins respectively), might impair efficient nuclear entry of the excess fusion protein product. In contrast, and consistently with the STED experiments, the SNAP-tag fusion allowed an efficient labeling with a clear synaptic enrichment comparable to the ones obtained with the other validated fusions (**Fig. 6**).

Altogether these results demonstrate that Xph15 and Xph20 constitute robust and valuable modules to engineer super-resolution imaging probes for endogenous PSD-95. Indeed, by adapting the recognition and the reporting modules together with the use of a system for regulation of the probe production, we show that they can be easily exploited to provide a straightforward access to both the nanoscale mapping and the dynamics of this key synaptic protein.

### Targeting sensors to synapses

With the series of fluorescent or self-labeling protein fusions to Xph15 and 20, we have demonstrated the efficiency of the evolved binders to be used as the targeting module to report on the localization of endogenous PSD-95. Considering the highly enriched distribution of PSD-95 at excitatory post-synapses, we sought to expand the scope of application of Xph15/20 by exploiting their binding properties to target sensors or bioactive proteins at the postsynapse.

To validate this strategy, we used the genetically encoded calcium reporter GCaMP (**Chen et al., 2013**; **Dana et al., 2019**) with the aim to generate a direct fluorescent indicator of individual synapse activity (**Fig. 7a**). A first attempt with GCaMP6f (**Chen et al., 2013**) as simple fusion to Xph15 expressed in primary culture neurons clearly indicated the feasibility of the approach (**Supplementary Fig. 9a and b, movies 1 and 2**). Indeed, even in the absence of the regulation system, a clear synaptic enrichment of the engineered calcium reporter was observed in comparison to the original sensor expressed alone. Expression levels were consistently low for the engineered construct, which can explain why the regulation system was not needed here to prevent excess probe production. We next attempted to improve the tool by using Xph20 as a stronger binder, GCaMP7f (**Dana et al., 2019**) as a brighter reporter, a stronger promoter (CAG instead of CMV), as well as the expression regulation system.

The two modified reporters (with and without the expression regulation system) were co-expressed with Homer-DsRed as a synaptic marker. Expression levels of the reporter were higher with the CAG promoter both in the absence and presence of the regulation system. However, in this case, the latter was necessary to allow a clear synaptic enrichment of GCaMP7f (**Fig. 7b, c and c**) as its absence, combined with higher expression levels, led to a homogenous distribution of the calcium reporter in the dendrite (**Supplementary Fig. 9**). Altogether, these results demonstrate that both the Xph15 and Xph20 binding modules can also be exploited to target gene-encoded

## Discussion

With the objective to develop imaging probes to monitor endogenous PSD-95, we have exploited a series of evolved synthetic binders of PSD-95 tandem PDZ domains derived from the ^10^FN3 domain. Taking advantage of both their unique paralog-specific recognition properties and their respective epitopes all situated on regions distant from the PDZ domains binding groove, we have first confirmed that binding to their target was not detectably affecting the PDZ domains nor the full-length protein function. Their use as ReMoRA endogenous PSD-95 probes in the form of fusions to fluorescent proteins was then evaluated in comparison to both antibodies and a similar -but not specific- monobody. The tools were next further engineered to adapt them for super-resolution imaging applications. We demonstrated that the resulting probes could be exploited with STED, PALM, and DNA-PAINT techniques to provide nanoscale mapping as well as dynamics information on endogenous PSD-95. Finally, we also showed that the binders can be employed to enrich active protein-based modules, such as calcium fluorescent reporters, at the excitatory post-synapse.

The three monobodies we considered in this study were all selected primarily based on their capacity to discriminate PSD-95 from other strongly homologous paralogs (PSD-93, SAP97, and SAP102). As shown before, this remarkable specificity results from their ability to recognize epitopes situated in regions distant from the targeted PDZ domains binding groove. Indeed, while the main site of interaction of the PDZ domains with their native protein partners is conserved across paralogs, their sequences are not strictly identical outside of these regions. Consequentially, binding module other than fluorescent proteins to excitatory synapses. In the case of the GCaMP reporter series, we also validate its compatibility with the gene regulation system in order to achieve a clear synaptic enrichment of the probe, of the PSD-95 specific monobodies does not obstruct the PDZ domain binding grooves. We show here, however, that while Xph15 and Xph20, which bind exclusively to the first PDZ domain, do not detectably affect the domain nor the full-length protein function, the situation is slightly different for Xph18. Indeed, this evolved ^10^FN3 domain presents an epitope that encompasses regions on both PDZ domains 1 and 2. As a result, binding of Xph18 locks the two domains, otherwise free to rotate around a short linker, in a specific conformation. This conformational constraint was only detectably detrimental to the interactions of synthetic divalent PDZ domain ligands. We therefore excluded this binder from imaging applications to avoid potential impact on PSD-95 activity, even if its expression does not seem to affect AMPAR stabilization at synapses not synaptic currents.

As previously reported, the Xph15 and Xph20 share very similar epitopes. Importantly, these epitopes are conserved in a number of species (e.g. rodents, human…) and are not subject to post-translational modifications. These features therefore guarantee a large spectrum of applications. Furthermore, we note that in the context of intrabody-like approaches, the ^10^FN3 scaffold, from which the binders are derived, is devoid of internal disulfide bonds, typically found for instance in antibody fragments, and thereby alleviating the risk of susceptibility to the intracellular reducing environment. Despite the differences in affinities and binding kinetics of Xph15 and Xph20, both allowed an efficient and specific labeling of PSD-95 independently of the associated reported group (eGFP, mNeonGreen, SNAP-tag, mEos3.2, GCaMP). Xph20, as the tightest binder, should therefore be preferred for most applications. However, the faster binding kinetics of Xph15 can also be exploited to favor rapid renewal of the probes in live conditions, a decisive advantage when photobleaching prevents time-based experiments.

With the growing access and interest for intrabodies or their synthetic equivalents, there is a need to develop strategies to adapt the expression level of the probe to its target, in particular in the case of imaging applications. We have opted here for a regulation system developed by the group of Don Arnold for a similar application (**Gross *et al.*, 2013**). It relies on the use of the excess (unbound) pool of probes to turn off further synthetic gene expression. In other words, the system is efficient if, on the one hand, the targeted protein is not nuclear and, on the other, the affinity of the evolved binder for the target is superior to the one of the appended zinc finger for its binding motif incorporated into the expression plasmid. Neuronal proteins that are located on cellular processes (dendrites or axons) are perfectly adapted for this strategy as the inevitable accumulation of fluorescent probes in the nucleus is not problematic for imaging purposes. We have observed here that the regulation system was functional for evolved binding modules with affinities in the 1-0.1 μM range in combination with a highly expressed target such as PSD-95. Indeed, for all probes and imaging techniques the expression profile was consistent with what would be expected from a directly labeled PSD-95 as confirmed by the strong co-localizations observed for the intrabodies and anti-PSD-95 antibody staining and the estimation of synaptic copy number by PALM. Furthermore, spt-PALM analysis revealed a major population of probes in a mildly diffusive state, as would be expected from PSD-95-bound reporters. However, we note that while most of the cargos we tried were compatible with this approach, the specific use of mScarlet-I and HaloTag resulted in the failure of the regulation system for reasons that are still unclear. The group that developed the regulation system has demonstrated that two orthogonal zinc finger systems could be used in concert (**Gross *et al.*, 2013**). Alternative methods to regulate the effective expression levels of the probe in tune with the one of its molecular targets would also be highly valuable for multiplexing applications as well as for systems (target, binder or cargo) outside of the optimal conditions mentioned above. Developing probes that undergo fast degradation unless bound to their target constitutes an interesting alternative that has been successfully used for the nanobody scaffold (**Gerdes *et al.*, 2020**; **Keller *et al.*, 2018**; **Tang *et al.*, 2016**). Another strategy for imaging applications would consist in conditioning the resulting fluorescence rather than the probe stability to its target binding by the development of fluorogenic probes (**Wongso *et al.*, 2017**).

We have demonstrated here that the probes could be adapted to comply with a number of fluorescence-based imaging techniques. Besides the advantages of the system to monitor endogenous PSD-95 in live or fixed conditions with standard imaging procedures, super-resolution imaging approaches can also be readily accessed with adequate probe engineering. Live imaging and protein dynamics investigations can be performed by exploiting STED or spt-PALM techniques. In the case of live STED, hour timescale studies will benefit from the straightforward use of most fluorescent protein fusions, whereas for studies that require a faster temporal resolution (minute timescale), coupling of brighter and more photorobust organic dyes can be achieved by the use of the SNAP-Tag. Precise nanoscale mapping of the protein target is accessed in fixed conditions either by STED with most probes, by PALM with photoswitchable fluorescent proteins such as mEos3.2, or by DNA-PAINT with a SNAP-Tag fusion as an anchoring point for the docking DNA strand. This large variety of imaging techniques applied to endogenous proteins highlights the potential of the labeling strategy compared to more conventional labeling schemes using either antibodies or direct genetic fusions. The strategy can be easily implemented to other imaging techniques, and for instance, STORM imaging could be achieved using either the SNAP-tag or the eGFP-based probes with respectively a BG or anti-GFP nanobody functionalized with dyes possessing photoswitching properties such as from the Cy5 cyanine family. Importantly, given the central role of PSD-95 in synaptic function, we anticipate that the probes will open up numerous possibilities for investigations against neuronal targets by providing straightforward solutions for the implementation of labeling/imaging strategies for multi-proteins studies.

As mentioned above, two other small synthetic PSD-95 binders have been recently reported by other groups in the context of imaging applications. One is a single-chain variable fragment (scFv), PF11, that was isolated against the palmitoylated form of PSD-95 and used as an intrabody (**Fukata *et al.*, 2013**). While the epitope was not clearly identified, the study showed recognition by the scFv of a conformational variant of PSD-95 that implied both N-terminal palmitoylation and the C-terminal part of PSD-95. Specificity was confirmed against PSD-93, one of the closest PSD-95 paralog that also possesses a palmitoylation site in its main isoform. The other binder is a monobody, therefore in the same synthetic binder scaffold class as Xph15 and Xph20, isolated from a selection performed against the C-terminal domains of PSD-95 (SH3 and guanylate kinase domains) (**Gross *et al.*, 2013**). The epitope was also not determined in this study, and the isolated monobody, PSD95.FingR, was shown to also recognize SAP97 and SAP102 paralogs but not PSD-93, a property that was exploited to investigate the role of SAP97/Dlg1 in cell polarity (**Li *et al.*, 2018**). In both cases affinities were not determined but the binders performed efficiently as intrabody-type probes for endogenous PSD-95. However, the absence of defined epitopes for both PF11 and PSD95.FingR does not allow to convincingly rule out possible perturbations of some of PSD-95 functions when any of the two probes is bound. PSD-95 is indeed a multidomain scaffold protein with a long list of identified partners (**Dosemeci *et al.*, 2007**; **Won *et al.*, 2017**; **Zhu *et al.*, 2020**) as well as numerous post-translational regulation sites (**Vallejo *et al.*, 2017**), which complicate evaluation of the impact resulting from a synthetic binder interaction. In addition, recent studies support the idea that the protein should not be viewed as a passive scaffolding element of the synapse but rather as an active actor with a capacity to respond to partners binding (**Rademacher *et al.*, 2019**; **Zeng *et al.*, 2018**). In this context, we note that the results we obtained with Xph18 illustrate the difficulty to establish with certainty whether a synthetic binder may impact the activity of its target even when the epitope is known. Indeed, while we could demonstrate that this particular monobody had a clear impact on PSD-95 conformation suggesting a plausible modification of its behavior in its native environment, we did not observe detectable perturbation of PSD-95 basic functions in basal conditions.

In comparison to PF11 and FingR.PSD-95, our study shows that Xph15 and Xph20 constitute valuable complementary molecular tools for standard imaging applications based on their unique specificity profile. They recognize both palmitoylated and non-palmitoylated PSD-95 and can discriminate PSD-95 vs its paralogs. Importantly, they present the net advantage of being characterized with respect to the identity of their respective epitope. While this was critical to clarify the molecular origin of the binders specificity for PSD-95, it also allowed us to relevantly adapt their evaluation in order to confirm the absence of impact of the probes on the target protein function. Critically, Xph15 and Xph20 remarkable specificity, as well as their binding properties, have allowed us to engineer the binders as super-resolution imaging probes to investigate endogenous PSD-95.

Besides the use of evolved synthetic binders as a strategy to label endogenous PSD-95 in live conditions, a number of genetic approaches have been reported. They all rely on gene editing methods and are typically used to generate PSD-95 fluorescent protein (**Broadhead *et al.*, 2016**; **Fortin *et al.*, 2014**; **Willems *et al.*, 2020**; **Zhu *et al.*, 2020**) or engineered self-labeling enzyme fusions (**Masch *et al.*, 2018**). Comparatively, their main advantages over expressed exogenous probes are the ideal stoichiometric labeling (one fluorophore per target protein, to be tempered by the notion of effective labeling efficiency of the fluorophore (**Thevathasan *et al.*, 2019**)) together, for the knock-in approaches, with the possibility to achieve global labeling.

In contrast, the ReMoRA or intrabody-based approaches benefit from their ease of implementation by relying on standard cell biology techniques for the genetically encoded probe delivery (transfection, electroporation or virus-mediated delivery). Indeed, gene editing methods are still not accessible in routine use to most laboratories and are also not amenable to downstream adaptation to various imaging techniques, the possibilities being imposed by the initial choice of the fluorescent module. The modular design of the synthetic binder-based probes provides in turn a system more adapted to engineering and optimization (binding module, fluorescent system, promoter…). Furthermore, we note that the rapid renewal of the probe obtained with the fast kinetics binders can be advantageous for imaging purposes over genetic modification of PSD-95 as its turn-over is slow in basal conditions.

In conclusion, we provide here a set of powerful probes for targeting PSD-95 with main applications for endogenous protein imaging as well as synaptic enrichment of active protein modules such as activity reporters. In comparison to other similar existing tools, the evolved monobodies described here constitute to this day the only binding modules displaying a strict specificity for PSD-95 regardless of its post-translational modification state. The molecular understanding of their mode of binding comforts our results, indicating undetectable perturbation of PSD-95 function. The probes presented here, which benefit from the simplicity of use of the ReMoRA design, provide direct access to different super-resolution imaging techniques. We anticipate that beyond the direct benefit for nanoscale mapping and (single molecule) dynamics investigations of endogenous PSD-95, these probes will turn invaluable for investigations that require the implementation of multiplexing imaging strategies.

## Materials and Methods

### Plasmid construction

The plasmids generated and the primers used in this study are listed in **Supplementary Tables 2 and 3**, respectively. Plasmids for protein production were previously described (**Rimbault *et al.*, 2019**; **Sainlos *et al.*, 2009**). Briefly, for bacterial expression, the first two PDZ domains of PSD-95 were subcloned into pET-NO to produce a N-terminal fusion with an octa-His tag and a TEV protease cleavage site. The Xph clones were subcloned into the pIGc vector to generate C-terminal fusions with a deca-His tag. For FRET experiments, PSD-95-eGFP and stargazin-mCherry were previously described (**Sainlos *et al.*, 2011**). Plasmids for soluble Xph clone expression were obtained by replacing the eGFP-CCR5 ZF-KRAB(A) fragment from the corresponding pCAG vector (gift from Don Arnold, USC, Addgene #46295) (**Gross *et al.*, 2013**) by an octa-His and HA tags using BglII and BsrGI restriction sites. Plasmid for soluble PSD-95 PDZ domain 2 expression was obtained by first replacing eGFP into pEGFP-N1 by mIRFP via BamHI and BsrGI restriction sites (gift from M Davidson, Florida State University, and X Shu, UCSF, Addgene #54620) (**Yu *et al.*, 2015**) and then subcloning the PDZ domain using HindIII and BamHI restriction sites. The PDZ domain-based FRET reporter was obtained as previously described (**Rimbault *et al.*, 2019**) but here without mutation of the first domain. The plasmid for expression of soluble Xph15 and 18 with a miRFP670 nuclear reporter were generated as described for the one with Xph20 (**Rimbault *et al.*, 2019**).

For imaging, Xph15, Xph18, and Xph20 were subcloned into pCAG_PSD95.FingR-eGFP-CCR5TC (gift from Don Arnold, USC, Addgene #46295) (**Gross *et al.*, 2013**) using KpnI and BglII restriction sites. Other fluorescent modules, mRuby2 (gift from Michael Lin, Addgene #40260) (**Lam *et al.*, 2012**), mScarlet-I (gift from Dorus Gadella, Addgene #98821) (**Bindels *et al.*, 2017**), mEos3.2 (gift from Michael Davidson & Tao Xu, Addgene 54525) (**Zhang *et al.*, 2012**), mNeonGreen (obtained by gene synthesis, Eurofins) (**Shaner *et al.*, 2013**), HaloTag (Promega, cat no G7971) and SNAPf (New England Biolabs, cat no N9183S) were next inserted in place of eGFP in the corresponding vector using BglII and NheI sites after an initial modification of the source vectors to introduce an NheI site between the fluorescent module and CCR5 ZF. GCaMP6f (**Chen *et al.*, 2013**) and GCaMP7f (**Dana *et al.*, 2019**) expressing plasmids were gifts from Douglas Kim & GENIE Project (Addgene #40755 and #104483 respectively). Xph15 was subcloned as a N-terminal fusion to GCaMP6f by using BglII and Sal I restriction sites and GCaMP7f was subcloned C-terminally to Xph20 into pCAG_Xph20-eGFP-CCR5TC using BglII and either MluI or NheI sites for removal or conservation of the eGFP-CCR5 ZF-KRAB(A) fragment, respectively.

### Protein production

Proteins were expressed and purified as previously described (**Rimbault *et al.*, 2019**). Briefly, His-tagged proteins were either produced in *E. coli* BL21 CodonPlus (DE3)-RIPL competent cells (Agilent, 230280) using auto-induction protocols (**Studier, 2005**) at 16 °C for 20 h or in BL21 pLysY (New England Biolabs, C3010I) for isotopically labelled proteins with IPTG induction for 16 h at 20 °C. Proteins were first isolated by IMAC using Ni-charged resins then further purified by size exclusion chromatography (SEC). An intermediate step of affinity tag removal by incubation with the TEV protease was added before the SEC step for isotopically-labelled proteins. The recovered proteins were concentrated, aliquoted and flash-frozen with liquid nitrogen for conservation at -80 °C.

### Peptide synthesis

Peptides were synthesized as previously described (**Rimbault *et al.*, 2019**). Briefly, amino acids were assembled at 0.05 mmol scale by automated solid-phase peptide synthesis on a CEM μwaves Liberty-1 synthesizer (Saclay, France) following standard coupling protocols. The divalent ligand [Stg_15_]_2_ was obtained by using copper-catalyzed click chemistry on resin harboring a mix of sequences functionalized by azide and alkyne groups as described previously (**Sainlos *et al.*, 2011**). Briefly, a 7:3.5 mixture of Fmoc-Lys(N_3_)-OH and pentynoic acid was manually coupled to the deprotected N-terminal amino group of elongated peptides on resin followed by copper(I)-catalyzed azide-alkyne cycloaddition in DMF/4-methylpiperidine (8:2) with CuI (5 eq), ascorbic acid (10 eq) and aminoguanidine (10 eq). N-free peptide resins were derivatized with acetyl groups or further elongated with a PEG linker (Fmoc-TTDSOH, 19 atoms, Iris Biotech, FAA1568) and fluorescein isothiocyanate. Peptides were purified by RP-HPLC with a semi-preparative column (YMC Ci8, ODS-A 5/120, 250 x 20 mm) and characterized by analytical RP-HPLC and MALDITOF. Peptides were lyophilized and stored at -80 °C until usage.

### NMR spectroscopy

NMR spectra were recorded at 298 K using a Bruker Avance III 700 MHz spectrometer equipped with a triple resonance gradient standard probe. Topspin version 4.1 (Bruker BioSpin) was used for data collection. Spectra processing used NMRPipe (**Delaglio *et al.*, 1995**). with analysis by using Sparky 3 (T. D. Goddard and D. G. Kneller, University of California). Titration of 200 μM ^15^N-labelled PSD-95 PDZ1-PDZ2 in PBS with a stock solution of 10 mM Stargazin C-terminal peptide (Ac-YSLHANTANRRTTPV) was followed by 1D ^1^H and 2D ^15^N-HSQC spectra. Titration points include 0, 40, 80, 120, 160, 200, 240, 280, 320, 360, 400 and 440 μM peptide, corresponding to 0.2, 0.4, 0.6, 0.8, 1, 1.2, 1.4, 1.6, 1.8, 2 and 2.2 molar equivalents peptide:protein. The titration was repeated by using pre-assembled 1:1 complexes of 200 μM ^15^N-labelled PSD-95 PDZ1-PDZ2 with a slight molar excess (240 μM) of natural abundance Xph15, Xph18, or Xph20. Amide ^1^H,^15^N chemical shift assignments of unbound and bound [^15^N]PSD-95-12 were previously reported (**Rimbault *et al.*, 2019**).

### Fluorescence polarization assay

For direct titrations, the fluorescein-labelled stargazin peptide (10 nM) was titrated against a range of increasing concentrations of the different recombinant PDZ domains in a 100 μl final volume. Fluorescence Polarization was measured in millipolarization units (mP) at an excitation wavelength of 485±5 nm and an emission wavelength of 520±5 nm using a POLARstar Omega (BMG Labtech) microplate reader. Titrations were conducted at least in duplicate and measured twice. To determine the corresponding affinities (apparent K_*D*_), curves were fitted using a nonlinear regression fit formula (**Chang *et al.*, 2011**) with GraphPad Prism v7.04 after normalizing the values of each protein series between the initial unbound and the saturating states.

For competitive titrations, experiments were designed such that the starting polarization value represents 75% of the maximal shift of the direct titrations. For the divalent stargazin ligand, PSD-95-12 was used at a concentration of 90 nM. Tandem PDZ domains, bound to the fluorescein-labelled stargazin divalent peptide (10 nM), were titrated against a range of increasing concentrations of acetylated stargazin divalent ligand in a 100 μl final volume, in the presence of 5 μM of Xph clones. For the monovalent stargazin ligand, PSD-95-12 (at a concentration of 20 μM), bound to the fluorescein-labelled stargazin monovalent peptide (50 nM), was titrated against a range of increasing concentrations of stargazin peptides in a 100 μl final volume, in the presence of 20 μM of Xph18. Titrations were conducted as above at least in duplicate and measured three times. To determine the corresponding inhibition constant (*K*_i_), curves were fitted using a competition formula (**Pazos *et al.*, 2011**) with GraphPad Prism v7.04 after normalizing the values of each protein series between the initial unbound and the saturating states.

### FRET/FLIM assays

FRET/FLIM assays were performed as previously described (**Rimbault *et al.*, 2019**). Briefly, COS-7 cells (ECACC-87021302) in DMEM medium supplemented with Glutamax and 10 % FBS were transfected using a 2:1 ratio X-treme GENE HP DNA transfection reagent (Roche) per μg of plasmid DNA with a total of 0.5 μg DNA per well. Experiments were performed after 24 h of expression. Coverslips were transferred into a ludin chamber filled with 1 ml fresh Tyrode’s buffer (20 mM Glucose, 20 mM HEPES, 120 mM NaCl, 3.5 mM KCl, 2 mM MgCl_2_, 2 mM CaCl_2_, pH 7.4, osmolarity around 300 mOsm.kg*^-1^*and pre-equilibrated in a CO_2_ incubator at 37 °C).

Experiments with full-length PSD-95 were performed using the time domain analysis (TCSPC) method with a Leica DMR TCS SP2 AOBS on an inverted stand (Leica Microsystems, Mannheim, Germany). The pulsed light source was a tunable Ti:Sapphire laser (Chameleon, Coherent Laser Group, Santa Clara, CA, USA) used at 900 nm and 80 MHz, providing a 13 ns temporal window for lifetime measurements. The system was equipped with the TCSPC from Becker and Hickl (Berlin, Germany) and fluorescence decay curves were obtained using single spot mode of SPCM software (Becker and Hickl).

Experiments with the PSD-95-12-derived FRET reporter system were performed using the frequency domain analysis (LIFA) method a Leica DMI6000 (Leica Microsystems, Wetzlar, Germany) equipped with a confocal Scanner Unit CSU-X1 (Yokogawa Electric Corporation, Tokyo, Japan). The FLIM measurements were done with the LIFA fluorescence lifetime attachment (Lambert Instrument, Roden, Netherlands), and images were analyzed with the manufacturer’s software LI-FLIM software.

Lifetimes were referenced to a 1 μM solution of fluorescein in Tris-HCl (pH 10) or a solution of erythrosin B (1 mg.ml^−1^) that was set at 4.00 ns lifetime (for fluorescein) or 0.086 ns (for erythrosin B). For competition experiments, only cells presenting a high level of expression of the competitor or control as measured by mIRFP670 fluorescence were taken into consideration.

### Cell culture

Rat Hipppocampal E18 culture neurons were prepared using a previously described protocol (**Penn *et al.*, 2017**) with the following modifications: neuron cultures were maintained in Neurobasal medium (cat. No. 12348017 ThermoFisher Scientic) supplemented with 2 mM Lglutamine (ThermoFisher Scientic Cat. No. 25030-024) and SM1 Neuronal Supplement (Cat. No. 05711 STEMCELL technologies).

### Gene delivery

For electrophysiology experiments, neurons were transfected with Xph15, Xph18, or Xph20 using Effectene kit (Qiagen N.V., Venlo, Netherlands) at 7-9 days in vitro (DIV). For immunostaining, FRAP, STED, and DNA-PAINT experiments, rat hippocampal neurons from E18 embryos were electroporated before plating with 1.5 μg of DNA using Nucleofector system (Lonza). For DNA-PAINT, primary hippocampal neurons were transfected using a standard calcium phosphate protocol at DIV 7-8 with Xph20-SNAP or Xph20-HaloTag and a soluble eGFP.

### Electrophysiology

Whole-cell patch clamp recordings were performed on Banker cultures of hippocampal neurons (13-17 DIV) expressing Xph15, Xph18, or Xp20 fused to eGFP. The experiments were carried out at room temperature in an extracellular solution (ECS) containing the following (in mM): 110 NaCl, 5.4 KCl, 10 HEPES, 10 glucose, 1.8 CaCl_2_, 0.8 MgCl_2_ (Sigma-Aldrich, St-Louis, USA); 250 mOsm; pH 7.4. To block voltage-gated sodium channels, 1 μM Tetrodotoxin (TTX; Tocris Bioscience, Bristol, UK) was added to the ECS. Intracellular solution (ICS) contained the following (in mM): 110 K-gluconate, 1.1 EGTA, 10 HEPES, 3 Na_2_ATP, 0.3 Na_2_GTP, 0.1 CaCl_2_, 5 MgCl_2_ (Sigma-Aldrich, St-Louis, USA); 240 mOsm; pH 7.2. Patch pipettes were pulled using a horizontal puller (P-97, Sutter Instrument) from borosilicate capillaries (GB150F-8P, Science Products GmbH) to resistances of 3-5 MΩ when filled with ICS. All recordings were performed using an EPC10 patch clamp amplifier operated with Patchmaster software (HEKA Elektronik). Data was acquired at 10 kHz and filtered at 3 kHz. Membrane capacitance was monitored frequently throughout the experiments and only cells with a series resistance <10 MΩ were analyzed.

Data were collected and stored on computer for off-line analysis using a software developed in-house (Detection Mini) to detect miniature synaptic events using a variable threshold. The amplitude and frequency of miniature excitatory postsynaptic currents (mEPSCs) were obtained for a minimum of 500 events.

Statistical values are given as mean ± SEM. Statistical significances were performed using GraphPad Prism software (San Diego, CA). Normally distributed data sets were tested by Student’s unpaired t-test for two independent groups.

### uPAINT

uPAINT was performed as previously reported (**Giannone *et al.*, 2010**) on dissociated neurons expressing Xph15, Xph18, or Xph20 fused to eGFP. Experiments took place at 13-16 DIV. Cells were imaged at 37 °C in an open chamber (Ludin chamber, Life Imaging Services, Switzerland) filled with 1 ml of Tyrode’s solution (in mM): 10 HEPES, 5 KCl, 100 NaCl, 2 MgCl*_2_*, 2 CaCl*_2_*, 15 glucose (pH 7.4). The chamber was mounted on an inverted microscope (Nikon Ti-Eclipse, Japan) equipped with a high 100X objective (1.49 NA), a TIRF device and an EMCCD camera (Evolve camera; Roper Scientific, Princeton Instruments, Trenton, NJ). Dendritic ROIs were selected based on eGFP signal. To track endogenous GluA2-containing AMPAR, an anti-GluA2 antibody given by E. Gouaux (Portland, OR) coupled to ATTO-647N (Atto-Tec, Siegen, Germany) was used. Stochastic labelling of the targeted protein by dye-coupled antibodies allowed the recording of thousands of trajectories lasting longer than 1 s. Recordings were made at 50 Hz using Metamorph software (Molecular Devices, USA), and analysis were performed with a homemade software developed under MetaMorph and kindly provided by J.B. Sibarita (Interdisciplinary Institute for Neuroscience).

### Immunostaining

At 23-27 DIV, Banker cultures expressing individual eGFP-tagged Xph or PSD95.FingR were stained with mouse monoclonal anti-PSD-95 (ThermoFischer Scientific cat. no. MA1-046). Briefly, neurons on coverslips were fixed 10 min using PFA 4%, washed with PBS, permeabilized with PBS-Triton-0.1% during 5 min, and washed again. After blocking with PBS-BSA 0.5%, neurons were stained with the PSD-95 antibody and after three washes with a secondary antibody (Goat anti-mouse Alexa 568, cat. no. A111031) for 45 min each. Neurons coverslips were mounted on Pro-Long Gold antifade reagent (ThermoFischer Scientific cat. no. P36934).

Images were acquired on a Leica DM5000 (Leica Microsystems, Wetzlar, Germany) with a HCX PL APO 63X oil NA 1.40 objective, a LED SOLA Light (Lumencor, Beaverton, USA) as fluorescence excitation source and a Flash4.0 V2 camera (Hamamatsu Photonics, Massy, France). Images quantifications were performed using tasks automatization with Metamorph. Following a background subtraction, the images were automatically thresholded to detect the positive objects for the synthetic binders (Xph15, Xph18, Xph20, or PSD95.FingR) and PSD-95. Enrichment was measured by the ratio between the fluorescence intensity of the positive objects for the synthetic binders and the shaft. Object colocalization was evaluated by determining regions of interest around positive objects for the synthetic binders and measuring the fluorescence intensity of these regions in the channel of PSD-95.

### FRAP

Banker cultures (21-23 DIV) in coverslips expressing eGFP fusions of Xph clones or full-length PSD-95 were mounted in a Ludin chamber (Life Imaging Services) and transferred to an inverted microscope (Leica, DMI 6000B) maintained at 37 °C. Fluorescence experiments were carried out in an extracellular solution containing (in mM): NaCl (120), KCl (3.5), MgCl_2_ (2), CaCl_2_ (2), D-glucose (10), HEPES (10) (pH 7.4, ∼270 mOsm), and transfected neurons were observed through a 63x oil objective (Leica, HC PL APO CS2, NA 1.4). GFP fluorescence was illuminated with 491 nm laser light using a high-speed spinning disk confocal scanner unit (Yokogawa CSU22-W1) and emission was captured with a sCMOS camera (Prime 95B, Photometrics). Microscope hardware was controlled with MetaMorph (Molecular Devices, v7.1.7) and ILAS2 system (Roper) softwares.

For FRAP experiments, the following protocol was used: 1) prebleaching (20 images at 3 s interval); 2) photobleaching of the regions of interest (10-15 ROIs per field of view, ROI=10 pixels, eq. to 2.3 μm), 3) fast recovery (40 images at 0.5 s interval) and 4) long-term recovery (10 mins recording at 3 s interval). For photobleaching, we used a 5 ms pulse of 488 nm laser light sufficient to reduce fluorescence by at least 50 %. Experiments where fluorescence dropped more than 20% in non-bleached regions during acquisition were discarded.

FRAP experiments were analyzed using an in-house developed macro to the ImageJ freeware (http://imagej.nih.gov/ij/). The source code is freely available from GitHub (https://github.com/fabricecordelieres/IJ-Macro_FRAP-MM), accompanied by a documentation and an example dataset. Briefly, as part of the macro, the FRAP region for each spine was imported from the Metamorph software to ImageJ’s ROI Manager using the Metamorph Companion plugin (https://github.com/fabricecordelieres/IJ-Plugin_Metamorph-Companion). From the data extracted by the macro, average intensity within the ROI was collected for each timepoint (F_*t*_), at first timepoint (pre-bleach, F_pb_) and immediately after the bleaching (F_0_). Simple normalization was performed as follows: FRAP_t_ =F_t_-F_ci_ /F_pb_-F_0_. The mean spine FRAP curve of each cell was subsequently fitted to a mono-exponential model using GraphPad Prism software.

### STED

Fixed neuronal cultures (DIV21) expressing GFP-tagged Xph20 were imaged with a glycerol immersion objective (Plan Apo 93x NA 1.3 motCORR). Cells were immunolabeled with MAP2 (Synaptic systems 188006 and anti-chicken AF594, ThermoFisher A11042) to identify dendritic draft. A 660-nm wavelength laser was used for GFP depletion. Acquisition parameters were: 20 nm pixel size, 4 times accumulated average per line and 200 Hz scan speed.

Banker cultures expressing mNeonGreen-tagged Xph15 or Xph20 were imaged at 37 °C in Tyrode’s solution. Live staining of rat hippocampal neurons transfected with both cytosolic eGFP and Xph15-SNAP-tag at DIV 10 was adapted from (**Bottanelli *et al.*, 2016**). Transfected neurons seeded on 18-mm coverslips at DIV 17 were incubated at 37 °C in the presence of 2 μl of aliquoted stock solution of the fluorescent ligand BG-SiR diluted in 250 μl of conditioned Neurobasal medium (final 5 μM BG-SiR). After 1 hr incubation, neurons were washed three times with 1 ml of CO_2_-equilibrated Neurobasal medium. Each wash was corresponding to a minimal 15-minutes incubation time with the fresh medium, to ensure removing all excess of unbound fluorescent ligand. Coverslip with neurons was then mounted in a Ludin chamber filled with 600 μl of pre-warmed Tyrode medium.

Live neurons were imaged with an inverted Leica SP8 STED microscope equipped with an oil immersion objective (Plan Apo 100X NA 1.4), white light laser 2 (WLL2, 470-670 nm, 80 MHz frequency, ca. 200 ps pulse duration), and internal hybrid detectors. A 775-nm pulsed wavelength laser (80 MHz frequency, ca. 600 ps pulse duration) was used to deplete SiR dye excited by the 647-nm laser line. To preserve neuron health, low STED power was used: time averaged measurements of STED laser power at the focal plane were showing a value lower than 20 mW (using S120C probe from Thorlabs). Other acquisition parameters were: 19 nm pixel size; 16 times average per line; bidirectional 400 Hz scan speed. Final images were processed in ImageJ as follow: gentle convolution using convolve plugin with a 3 x 3 kernel (1 1 1, 1 10 1, 1 1 1), slight chromatic correction to align GFP image with STED capture. Gamma correction of 0.5 was applied on neuron large view image to help seeing small synapses stained with SiR.

### (spt)PALM

Live or fixed (PFA 4%) cells were mounted in a Ludin chamber filled with 1 ml of Tyrode’s solution (in mM): 10 HEPES, 5 KCl, 100 NaCl, 2 MgCl_2_, 2 CaCl_2_, 15 glucose (pH7.4), and imaged at 37 °C. An inverted microscope (DMi8, Leica, Germany) equipped with a TIRF objective (160x 1.43 NA Leica, Germany), a Ilas^2^ TIRF device, and an Evolve EMCCD camera (Roper Scientific, Evry, France) was used for (spt)PALM recordings. Neurons expressing mEos3.2-tagged constructs were photo-activated using a 405-nm laser and the resulting photo-converted single-molecule fluorescence signal was excited with a 561-nm laser. The power of the 405-nm laser was adjusted to keep the number of the stochastically activated molecules constant and well separated during the acquisition. Images were acquired by image streaming for up to 4000 frames (sptPALM) or up to 20000 frames (PALM) at frame rate of 50 Hz using Metamorph software (Molecular Devices, USA), and analysis were performed with a homemade software developed under MetaMorph and kindly provided by J.B. Sibarita (Interdisciplinary Institute for Neuroscience).

### SMLM analysis

Localization and tracking reconnection of ATTO-647N (uPAINT) or mEos3.2 (PALM) signals were performed using homemade software developed as a MetaMorph plugin and kindly provided by J.B. Sibarita (Interdisciplinary Institute for Neuroscience) (**Kechkar *et al.*, 2013**). Single-molecule fluorescence could be identified by occurrence of fluorescence in the red channel and the defined minimum duration of fluorescence. Trajectories were reconstructed by a simulated annealing algorithm (**Racine *et al.*, 2006**), taking into account molecule localization and total intensity. Diffusion coefficients were calculated by linear fit of the first four points of the Mean Square Displacement plots.

PALM clusters analysis was performed using SR-Tesseler software as previously described (**Levet *et al.*, 2015b**).

### DNA-PAINT

Primary hippocampal neurons transfected with Xph20-SNAP and cytosolic eGFP were fixed at DIV 14-16 with 4% PFA in PBS for 20 min. Neurons were then quenched with 150 mM Glycine for 20 minutes, followed by simultaneous blocking and permeabilization for 90 min in PBS supplemented with 0.2% Triton-X-100 and 3% BSA. For SNAP-labeling, cells were incubated with 1 *μM* of SNAP-ligand-modified DNA oligomer in PBS supplemented with 0.5% BSA and 1 mM DTT for 1 hour.

Neurons were imaged at 25 °C in a Ludin chamber with an inverted motorized microscope (Nikon Ti) equipped with a CFI Apo TIRF 100x oil, NA 1.49 objective and a perfect focus system PFS-2, allowing long acquisition in TIRF illumination mode. For DNA-PAINT nanoscopy, neurons expressing Xph20-SNAP were first incubated for 15 min with 90 nm Gold Nanoparticles (Cytodiagnostics) to serve as fiducial markers. Xph20-SNAP was then visualized with Cy3b-labelled DNA imager strands, added to the Ludin chamber at variable concentrations (2-5 nM), as previously described (**Schnitzbauer *et al.*, 2017**). Cy3B-labelled strands were visualized with a 561 nm laser (Cobolt Jive). Fluorescence was collected by the combination of a dichroic and emission filters (dichroic: Di01-R561, emission: FF01-617/73, Semrock) and a sensitive sCMOS (scientific CMOS, ORCA-Flash4.0, Hammatasu). The acquisition was steered by Metamorph software (Molecular Devices) in streaming mode at 6.7 Hz. GFP was imaged using a conventional GFP filter cube (excitation: FF01-472/30, dichroic: FF-495Di02, emission: FF02-520/28, Semrock). Super-resolution DNA-PAINT reconstruction and drift correction were carried out as described before, using the software package Picasso (**Schnitzbauer *et al.*, 2017**).

### Calcium signaling imaging

Imaging of GCaMP6f and Xph-GCaMP6f was carried out in rat hippocampal dissociated cultures nucleoporated with the appropriate DNA on the day of the culture. Neurons were imaged at 13 to 18 DIV using a Nikon inverted microscope (Ti Eclipse) with an EMCCD camera (Evolve 512, Photometrics) controlled by Metamorph software (Molecular Devices) and equipped with a 60x/1.49 N.A. oil-immersion objective (Nikon). Images were acquired at a rate of ∼50 Hz. The imaging chamber (Ludin Chamber, Life Imaging Services) was perfused with extracellular buffer containing (in mM): 130 NaCl, 2.5 KCl, 3 CaCl_2_, 0.1 MgCl_2_, 10 glucose, 10 HEPES, 0.001 TTX, 0.05 PTX, (pH adjusted to 7.4 with NaOH and osmolarity adjusted to 280 mOsm) at room temperature. The fluorophores were excited with 488-nm laser lines, and imaged with the appropriate filters.

E18 rat hippocampal neurons were electroporated with GCaMP7f or Xph20-GCaMP7f and Homer1c-Dsred using the 4D Nucleofection system (Lonza) at DIV0, seeded on 18 mm glass coverslips, and cultured for 15 days. Imaging was performed by placing coverslips in a Ludin observation chamber (Life Imaging Services) in Mg^2^+-free Tyrode’s solution (15 mM D-glucose, 108 mM NaCl, 5 mM KCl, 2 mM CaCl2 and 25 mM HEPES, pH 7.4) containing 20 μM glycine inside a thermostatic chamber (37 °C) placed on an inverted microscope (Nikon Ti-E Eclipse) equipped with an EMCCD camera (Evolve 512, Photometrics) controlled by Metamorph software (Molecular Devices) and equipped with a 60x/1.49 N.A. oil-immersion objective (Nikon). Fluorescence was collected using a mercury lamp (Nikon Xcite) and appropriate filter sets (SemROCK).

For fig 7d. Quantification of synaptic enrichment was performed by segmenting Homer1c-DsRed clusters (synapses). These regions were transferred onto the GCaMP7f signal, and average intensity of GCaMP7f in these synaptic regions was measured and divided by the average intensity of the shaft area containing no homer-positive signal.

## Acknowledgments

This research was financially supported by grants from the Centre National de la Recherche Scientifique, the Conseil Régional d’Aquitaine, the France Biolmaging national infrastructure (grant ANR-10-INBS-04), the Agence Nationale de la Recherche (CheMoPPI, ANR-13-BS07-0019-01, OptoXL, ANR-16-CE16-0026) to C.P and M.S, the European Research Council to D.C., the Labex BRAIN (ANR-10-LABX-43) to C.R., Fondation pour la Recherche Médicale fellowship to B.C. We also thank the IINS cell culture facility and Emeline Verdier for technical assistance, the Biochemistry and Biophysics Core Facility of the Bordeaux Neurocampus both funded by the Labex BRAIN (ANR-10-LABX-43) and Y. Ruffin for technical assistance. Financial support from the IR-RMN-THC Fr3050 CNRS for conducting the research is gratefully acknowledged. We are grateful to M. Goillandeau for the upgrade of the mini analysis software, Dolors Grillo-Bosch for peptide synthesis, Kashyap Maruthi for making protein samples for the NMR studies, and Fabrice Cordelières for the FRAP analysis macro development.

## Author contributions

C.R. and M.S. designed the research and wrote the article with input from all authors. C.R. performed the biophysical experiments and generated the constructs with the help of C.B., C.G., I.G. C.D.M. designed NMR experiments, performed all NMR experiments and analyzed data. C.B., V.P. and C.P. performed the FRET/FLIM experiments. C.B. performed the immunostaining and initial imaging experiments. C.P. performed the analysis. E.T. and E.H performed the electrophysiology experiments. B.C. performed the uPAINT and PALM/sptPALM experiments. M.F.M. performed the FRAP experiments. C.P., M.F.M. and P.M. performed the STED experiments. J.F.N.V. performed the DNA-PAINT experiments with the supervision of G.G. E.T. and I.C. performed the GCaMP experiments. M.S. coordinated and oversaw the research project. All authors discussed the results and commented on the manuscript.

