## Supplementary Information for "Engineering paralog-specific PSD-95 synthetic binders as potent and minimally invasive imaging probes"

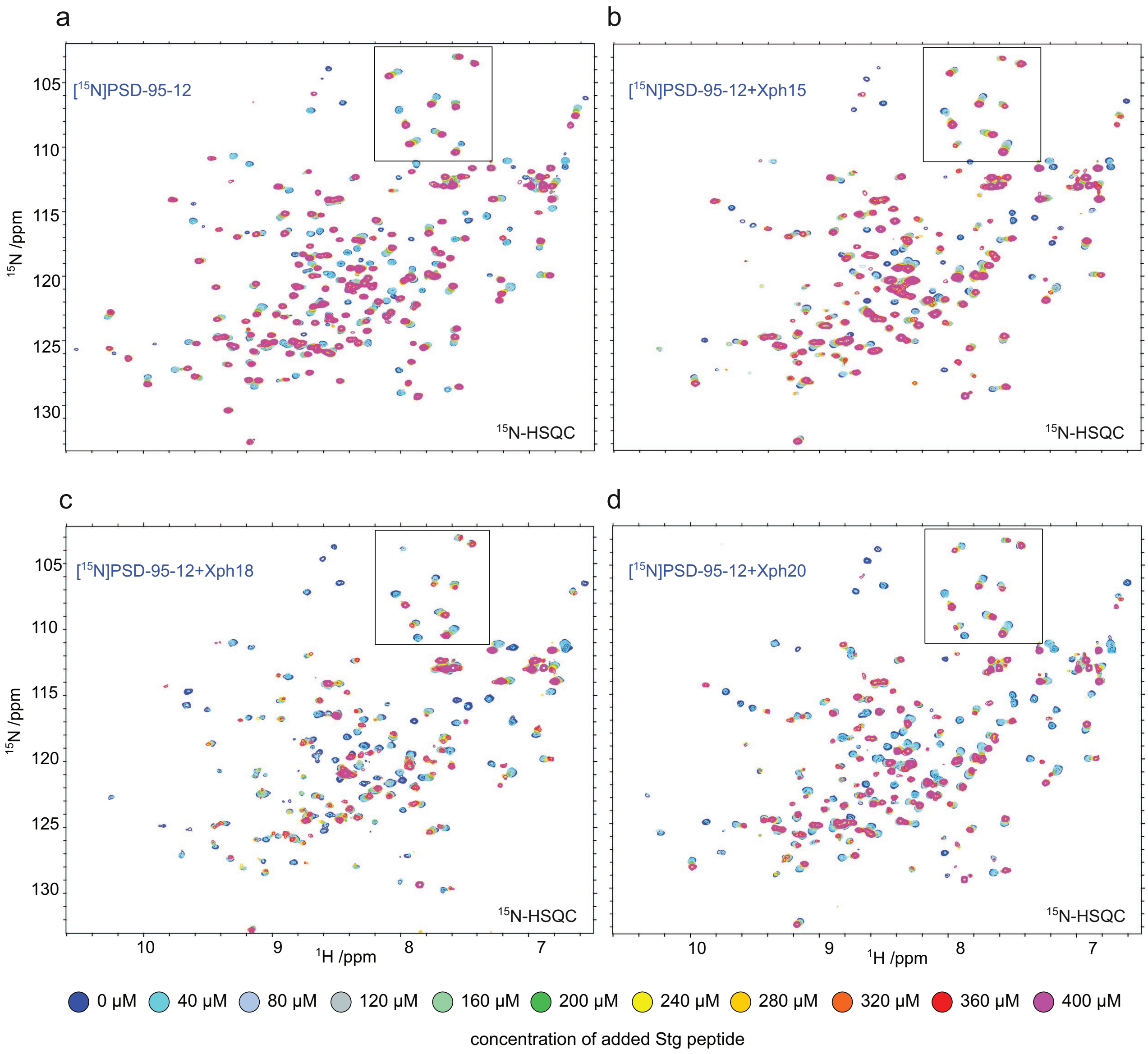


**Supplementary Figure 1 |** [$^{15}$N]-HSQC spectra collected on 200 µM [$^{15}$N]PSD-95-12 and titrated in with increasing concentrations of a monovalent stargazin-derived peptide (Stg) **(a)** in absence of binder, or in the presence of 240 µM Xph15 **(b)**, Xph18 **(c)**, or Xph20 **(d)**. The region in a box corresponds to the selected regions presented in **Fig 1b**.

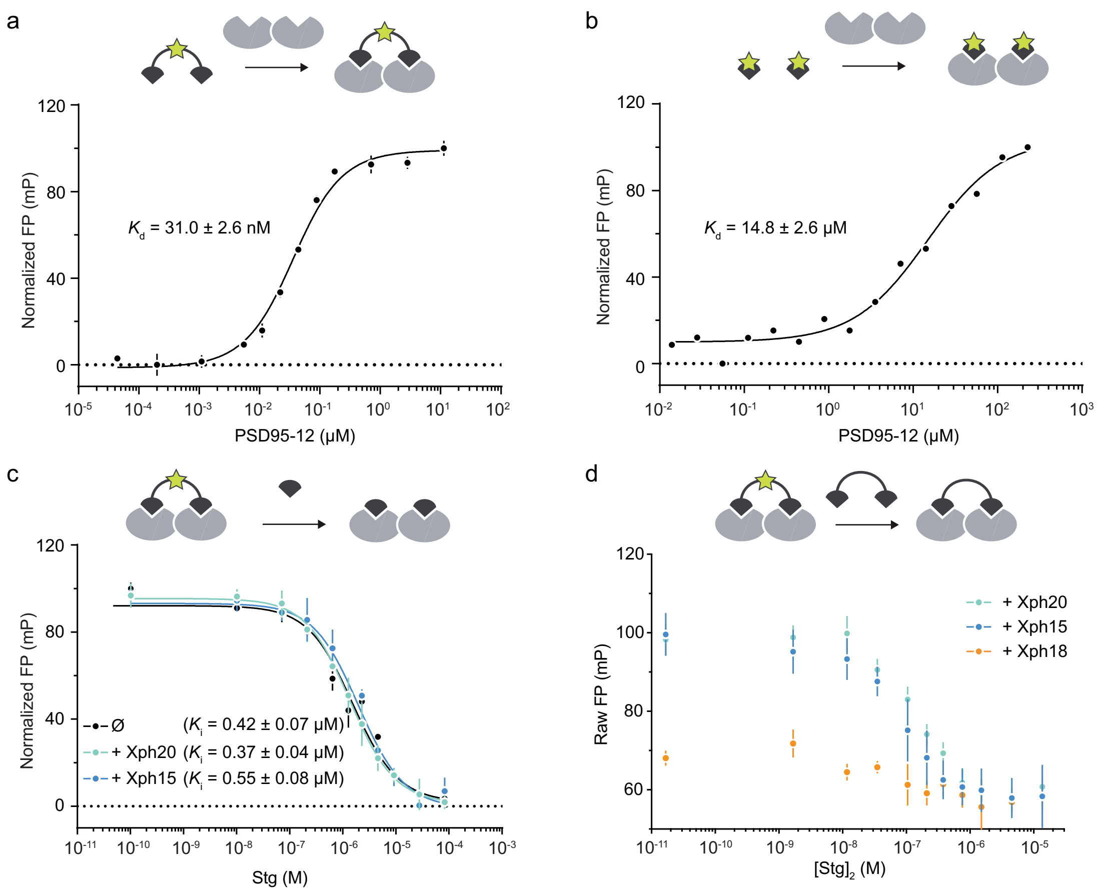


**Supplementary Figure 2 |** Fluorescence polarization titrations. **(a)** Direct titration of FITC-derived stargazin divalent ligand (10 nM) against PSD-95 tandem PDZ domains. Each data point represents the average of three measurements ± SD. The dissociation constants obtained by fitting are reported with the calculated SEM. **(b)** Direct titration of FITC-derived stargazin monovalent ligand (10 nM) against PSD-95 tandem PDZ domains. Each data point represents a single measurement. The dissociation constants obtained by fitting are reported with the calculated SEM. **(c)** Competitive titrations with non-fluorescent (acetylated) stargazin monovalent ligand against the complexes between FITC-derived stargazin divalent ligand, PSD-95 PDZ domain 1 and 2 and either Xph15, Xph20, or no binder (∅). Each data point represents the average of two independent measurements ± SD. The inhibition constants obtained by fitting are reported with the calculated SEM. **(d)** Competitive titrations with non-fluorescent (acetylated) stargazin divalent ligand against the complexes between FITC-derived stargazin divalent ligand, PSD-95 PDZ domain 1 and 2 and either Xph15, Xph20, or Xph18. Each data point represents the average of two independent measurements ± SD. The FP values are not normalized and indicate, in the presence of Xph18, an impaired binding of the fluorescent probe.

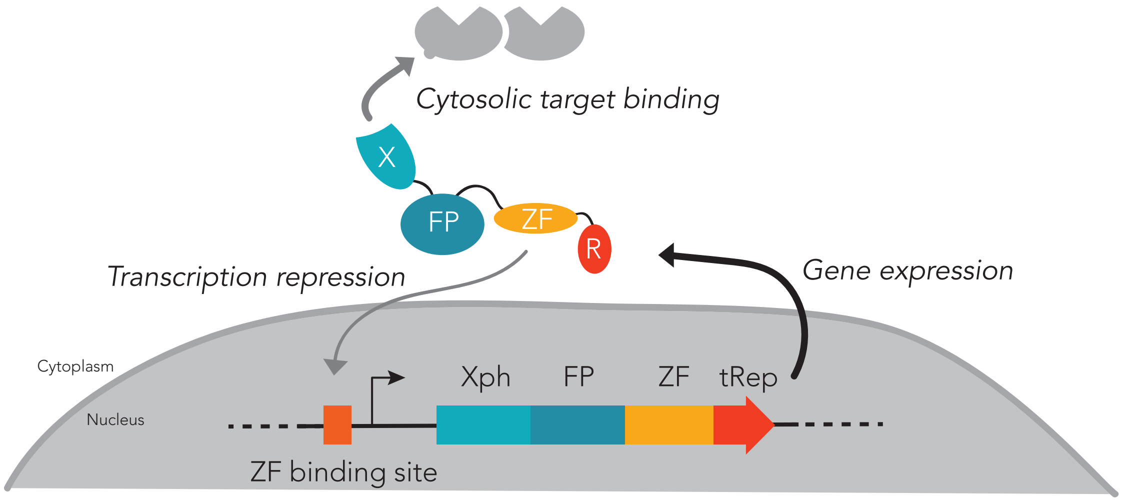


**Supplementary Figure 3 |** Expression regulation system. The system relies on a competition for the expressed probe between binding to its cytosolic target (favored) or preventing further transcription (unfavored until the target is saturated). Xph/X: binding module; FP: fluorescent protein; ZF: zinc finger; tRep/R: transcription repressor. For clarity, the expression regulation system is omitted in the schematic representation of the various probes used in neurons when its presence is not a variable.

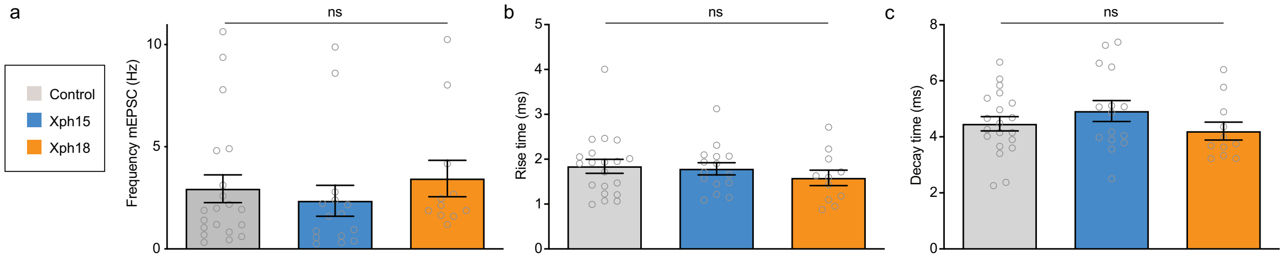


**Supplementary Figure 4 |** Spontaneous miniature excitatory postsynaptic currents properties based on the analysis of the 100 first events of control non-transfected or Xph15 and Xph18 (fused to eGFP and the expression regulation system) transfected culture neurons. **(a)** Frequency (in Hz, control: 2.95 ± 0.68 (n = 20); Xph15: 2.36 ± 0.75 (n = 15); Xph18: 3.44 ± 0.90 (n = 11); mean ± SEM with P>0.65 by ordinary one-way ANOVA). **(b)** Rise time (in ms, control: 1.84 ± 0.16 (n = 20); Xph15: 1.79 ± 0.14 (n = 15); Xph18: 1.58 ± 0.17 (n = 11); mean ± SEM with P>0.53 by ordinary one-way ANOVA). **(c)** Decay time (in ms, control: 4.46 ± 0.25 (n = 20); Xph15: 4.92 ± 0.37 (n = 15); Xph18: 4.20 ± 0.32 (n = 11); mean ± SEM with P>0.32 by ordinary one-way ANOVA).

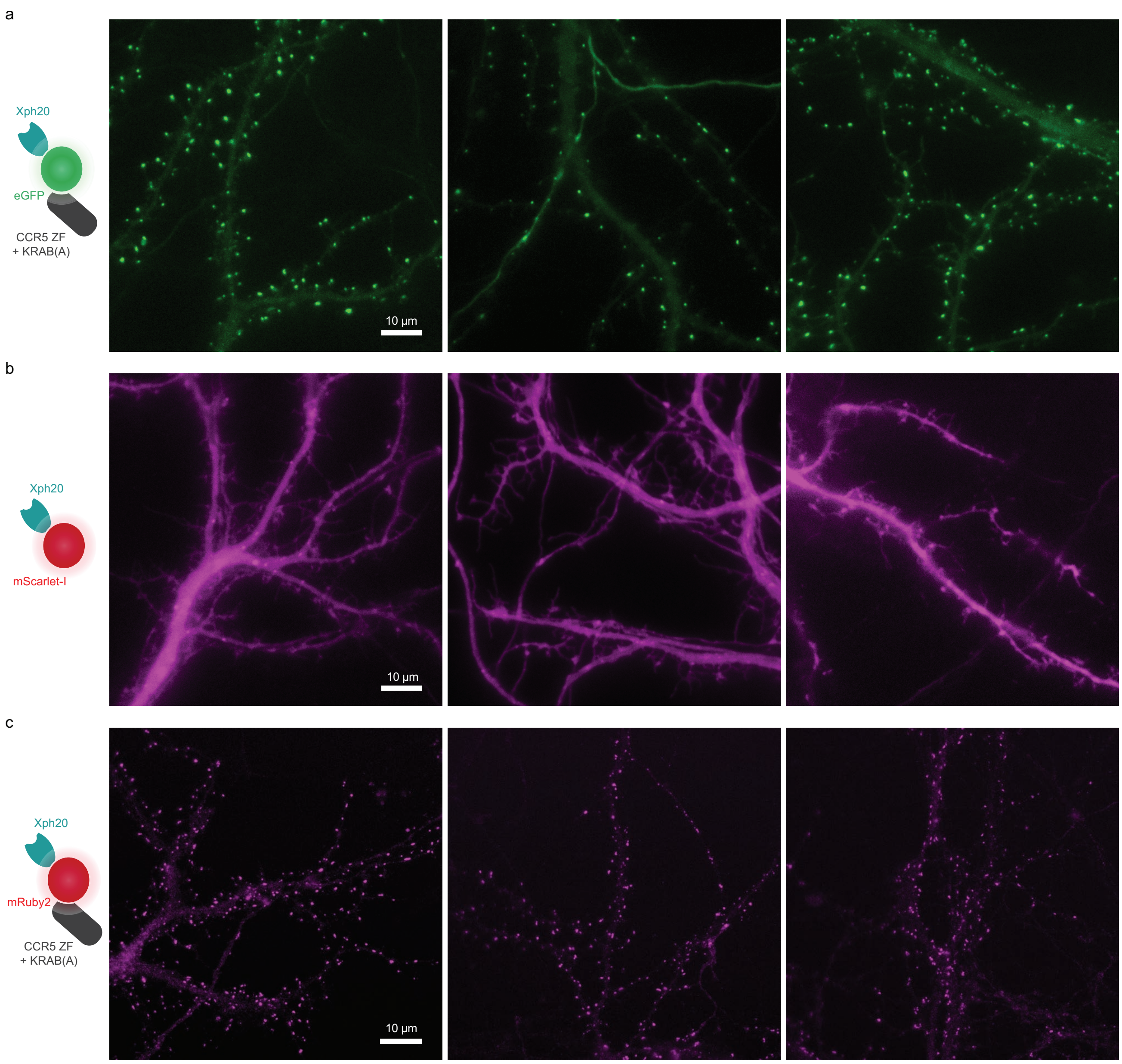


**Supplementary Figure 5 |** Regulated vs non-regulated expression of Xph20-FP fusion. **(a)** Representative images obtained on neurons transfected with Xph20-eGFP fusion (regulated with a zinc finger and a transcription repressor fusion together with the corresponding zinc finger binding site upstream of the promoter on the expression plasmid). **(b)** Representative images obtained on neurons from the same dissected as **(a)**, and transfected with Xph20-mScarlet-I fusion (non-regulated). **(c)** Representative images obtained on neurons transfected with Xph20-Ruby2 fusion (regulated).

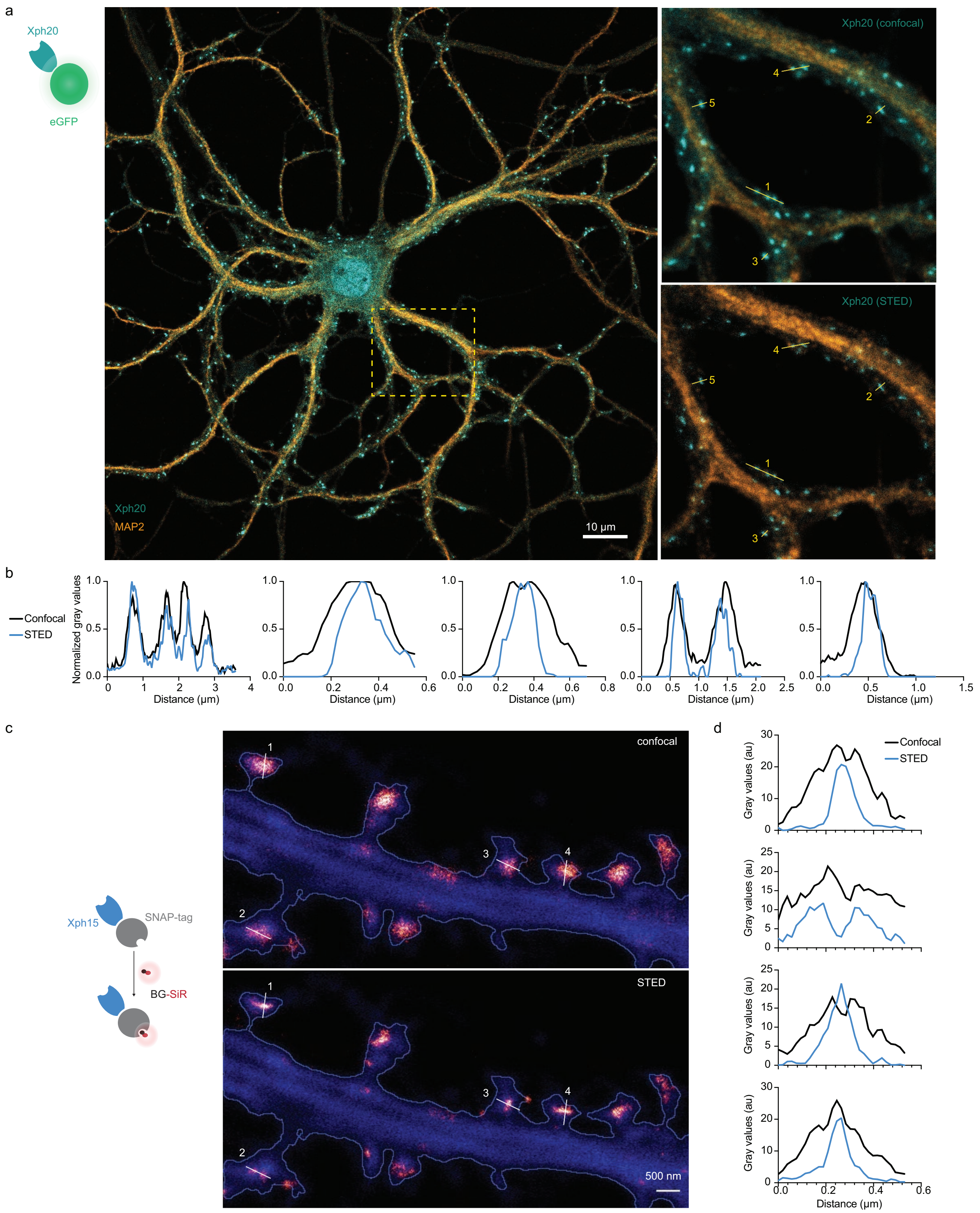


**Supplementary Figure 6 |** STED imaging **(a)** Representative image obtained on fixed neurons transfected with Xph20-eGFP fusion and immunolabeled with an antibody against MAP2 to identify dendritic shaft. Zoom region obtained by confocal and STED modalities. **(b)** Intensity profiles of Xph20-eGFP from the linescans indicated in the zoom region from **(a)**. **(c)** Representative images obtained by confocal and STED modalities on live neurons transfected with Xph15-SNAP-tag fusion and cytosolic GFP (blue), labeled with BG-SiR. **(d)** Intensity profiles of Xph15-SNAP-tag from the linescans indicated in the zoom region from **(c)**.

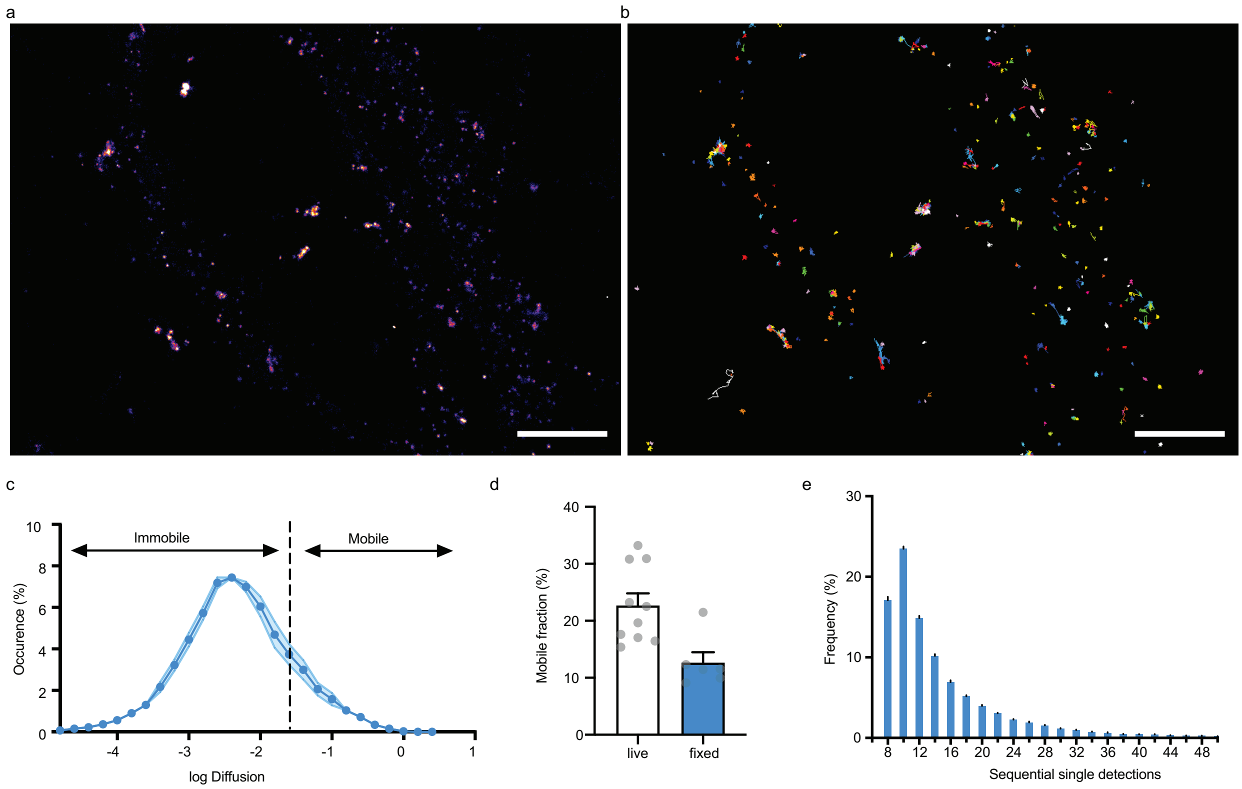


**Supplementary Figure 7 |** Evaluation of mEos3.2-derived probes for PALM applications. **a,b** Representative spt-PALM images of fixed culture neurons expressing Xph20-mEos3.2.). Super-resolution intensity map obtained by sptPALM from a sequence of 4,000 images of sparse single molecules of the photoconverted red from of mEos3.2 **(a)** and trajectories of Xph20-mEos3.2 **(b)**. **(c)** Average distribution of instantaneous diffusion coefficients obtained by spt-PALM of Xph20-mEos3.2. Error bars indicate cell-to-cell variability. **(d)** Percentage of the mobile fraction of probes (mean ± SEM, each dot represents a single cell). **(e)** Frequency distribution of sequential single Xph20-mEos3.2 detections.

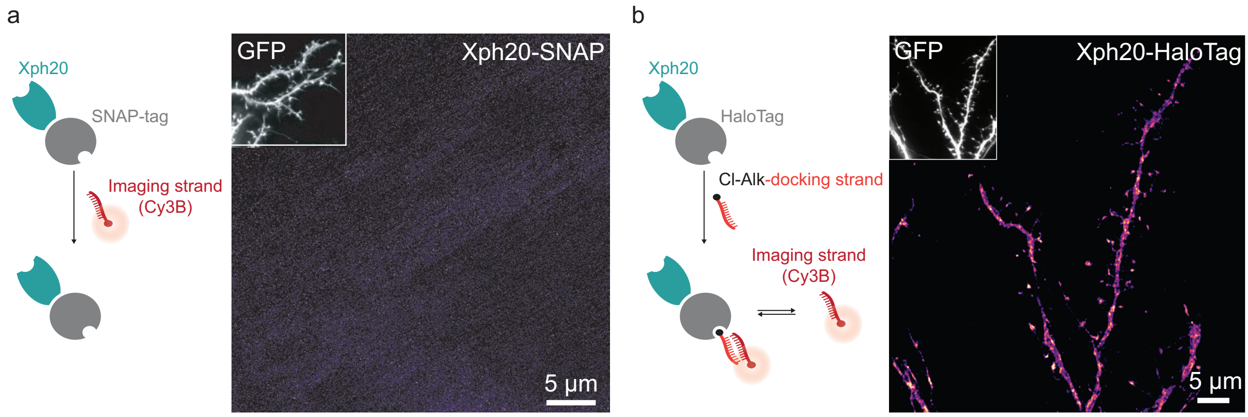


**Supplementary Figure 8 |** DNA-PAINT imaging**. (a)** Representative DNA-PAINT image obtained on neurons co-transfected with a soluble GFP marker and Xph20-SNAP-tag, not incubated with BG-docking strand and in the presence of Cy3-imaging strand. **(b)** Representative DNA-PAINT image obtained on neurons co-transfected with a soluble GFP marker and Xph20-HaloTag (Cl-Alk: chloroalkane). The reconstructed image shows an evident lack of synaptic enrichment

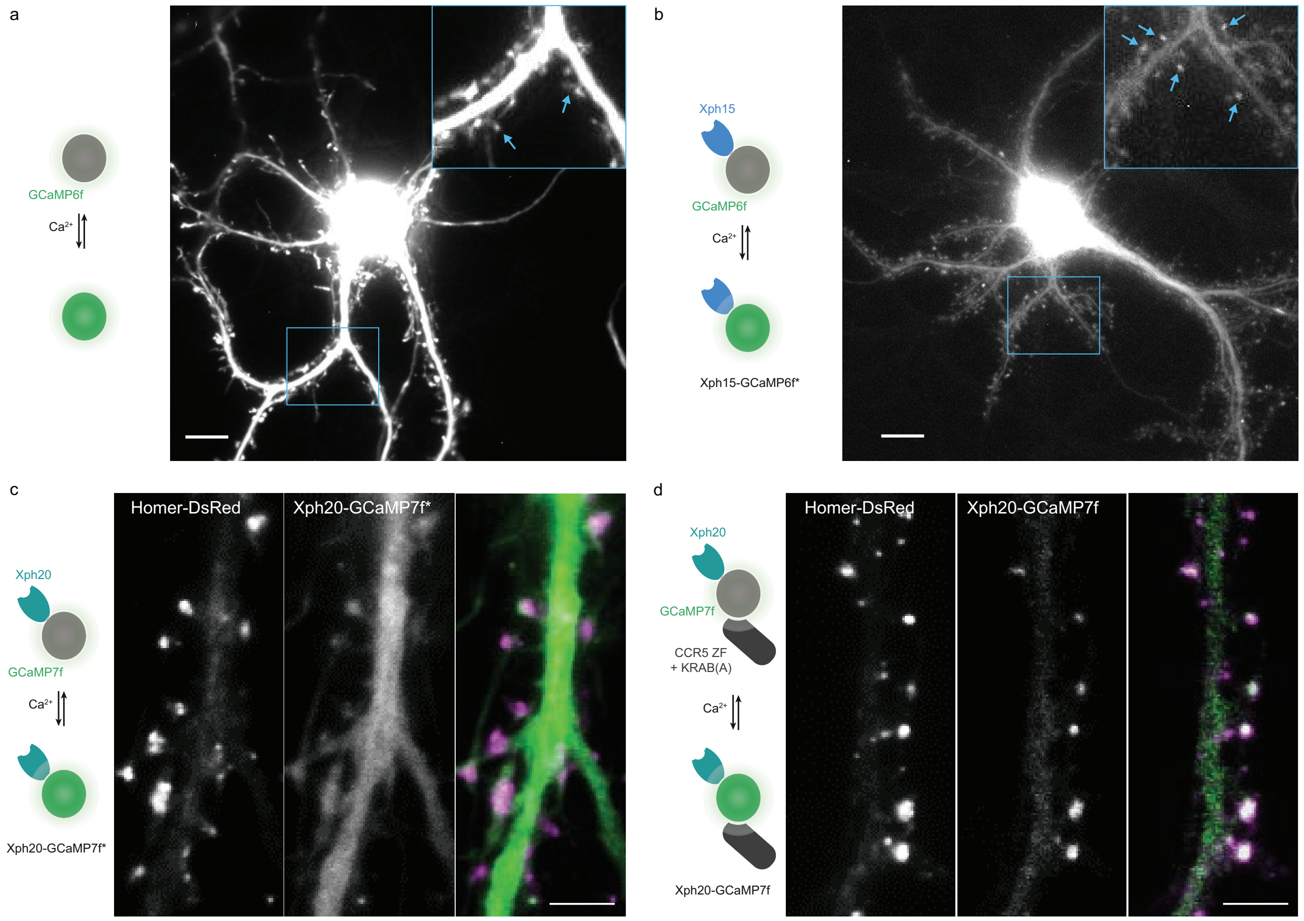


**Supplementary Figure 9 | (a)** and **(b)** Maximum projection of the fluorescence movies collected for hippocampal neurons expressing GCaMP6f **(a)** or Xph15-GCaMP6f* (non-regulated) **(b)**. In the insets, the blue arrows indicate individual spines. Scale bars, 10 μm. **(c)** and **(d)** Comparison of the expression profile of regulated (Xph20-GCaMP7f) vs non-regulated probe (Xph20-GCaMP7f*) for GCaMP7f synaptic targeting. **(c)** Representative images obtained on neurons co-transfected with a Homer-DsRed as a synaptic marker and GCaMP7f fused to Xph20 without the expression regulation system (Xph20-GCaMP7f*). **(d)** Representative images obtained on neurons co-transfected with a Homer-DsRed as a synaptic marker and GCaMP7f fused to Xph20 with the expression regulation system (Xph20-GCaMP7f). The expression regulation system allows to obtain a robust synaptic enrichment. Scale bars, 5 μm.

**Supplementary Table 1 |** FRAP data

| | Xph15 | Xph18 | Xph20 | PSD-95-EGFP |
|---|---|---|---|---|
| Plateau ± SEM (normalized) | 0.815 ± 0.005 | 0.707 ± 0.005 | 0.802 ± 0.005 | 0.462 ± 0.007 |
| Rate constant ± SEM ($s^{-1}$) | 0.073 ± 0.007 | 0.010 ± 0.001 | 0.011± 0.001 | 0.006 ± 0.001 |

**Supplementary Table 2 |** List of plasmids used in this work (c = commercial source; black dot = from this study).

| Entry | Plasmid name | Plasmid backbone | Origin of replication | Promoter | Gene | Purpose | Tag(s) | Figure(s) | Source ref | Addgene ID |
|---|---|---|---|---|---|---|---|---|---|---|
| p01 | NO-PSD-95-12 | pET-NO | pBR322 | T7 | PSD-95-12 [61-249] | NMR, FP | $His_8$; $TEV_{cs}$ | 1b,c,d; S1; S2 | 1 | — |
| p02 | pET-24a(+) | pET-24a(+) | pBR322 | T7 | Ø | Molecular biology | T7 tag; $His_6$ | — | c | — |
| p03 | pIGc-Xph20 | pIGc | pBR322 | T7 | Xph20 [S63K] | NMR, FP | $His_{10}$ | 1b,c; S1; S2c,d | 1 | 133030 |
| p04 | pIGc-Xph18 | pIGc | pBR322 | T7 | Xph18 [S63K] | NMR, FP | $His_{10}$ | 1b,d; S1; S2d | 1 | 133029 |
| p05 | pIGc-Xph15 | pIGc | pBR322 | T7 | Xph15 [S63K] | NMR, FP | $His_{10}$ | 1b,c; S1; S2c,d | 1 | 133028 |
| p06 | pIGc-Xph0 | pIGc | pBR322 | T7 | Xph0 | NMR, FP | $His_{10}$ | 1c | * | — |
| p07 | Stargazin mCherry [-21] | pcDNA3 | SV40 | CMV | Stargazin | FRET | mCherry at position -21 | 1e | 2 | — |
| p08 | PSD-95 eGFP | pcDNA3 | SV40 | CMV | PSD-95 | FRET | eGFP at position +253 | 1e; 3e,f,g | 2 | — |
| p09 | pCAG_Xph15 | pCAG | SV40 | CAG | Xph15 | FRET | HA | 1e | * | — |
| p10 | pCAG_Xph18 | pCAG | SV40 | CAG | Xph18 | FRET | HA | 1e | * | — |
| p11 | pCAG_Xph20 | pCAG | SV40 | CAG | Xph20 | FRET | HA | 1e | * | — |
| p12 | pCAG_Xph0 | pCAG | SV40 | CAG | Xph0 | FRET | HA | 1e | * | — |
| p13 | pmIFP-PSD-95-2 | pEGFP-N1 | SV40 | CMV | PSD-95-2 [155-249] | FRET | mIFP | 1e | * | — |
| p14 | pcDNA-FRET-PSD-95-12 (no stg) | pcDNA3 | SV40 | CMV | FRET sensor OFF ( negative control) | FRET | eGFP, mCherry | 1f | * | — |
| p15 | pcDNA-FRET-PSD-95-12 | pcDNA3 | SV40 | CMV | FRET sensor | FRET | eGFP, mCherry | 1f | * | — |
| p16 | pCAG-mIRFP670nuc-TEV-Xph15 | pCAG | SV40 | CAG | Xph15 [S63K], mIRFP670-Nuc | FRET | — | 1f | * | — |
| p17 | pCAG-mIRFP670nuc-TEV-Xph18 | pCAG | SV40 | CAG | Xph18 [S63K], mIRFP670-Nuc | FRET | — | 1f | * | — |
| p18 | pCAG-mIRFP670nuc-TEV-Xph20 | pCAG | SV40 | CAG | Xph20 [S63K], mIRFP670-Nuc | FRET | — | 1f | 1 | — |
| p19 | pCAG_Xph15-eGFP-CCR5TC | pCAG | SV40 | CAG | Xph15 - eGFP-CCR5 ZF-KRAB(A) | Imaging | eGFP | 2; 3; S4 | * | 135528 |
| p20 | pCAG_Xph18-eGFP-CCR5TC | pCAG | SV40 | CAG | Xph18 - eGFP-CCR5 ZF-KRAB(A) | Imaging | eGFP | 2; 3; S4 | * | 135529 |
| p21 | pCAG_Xph20-eGFP-CCR5TC | pCAG | SV40 | CAG | Xph20 - eGFP-CCR5 ZF-KRAB(A) | Imaging | eGFP | 2; 3; S5; S6a,b | * | 135530 |
| p22 | pCAG_PSD95.FingR-eGFP-CCR5TC | pCAG | SV40 | CAG | PSD95.FingR - eGFP-CCR5 ZF-KRAB(A) | Imaging | eGFP | 3 | 3 | 46295 |
| p23 | pCAG_Xph20-mRuby2-CCR5TC | pCAG | SV40 | CAG | Xph20 - mRuby2-CCR5 ZF-KRAB(A) | Imaging | mRuby2 | S5 | * | 135531 |
| p24 | pCAG_Xph20-mScarlet-I-CCR5TC | pCAG | SV40 | CAG | Xph20 - mScarlet-I-CCR5 ZF-KRAB(A) | Imaging | mScarlet-I | — | * | — |
| p25 | pTriex_Xph20-mScarlet-I | pTriex-5 | ColE1 | CMV | Xph20 - mScarlet-I-CCR5 ZF-KRAB(A) | Imaging | mScarlet-I | S5 | * | — |
| p26 | pCAG_Xph15-mNeonGreen-CCR5TC | pCAG | SV40 | CAG | Xph15 - mNeonGreen-CCR5 ZF-KRAB(A) | Imaging | mNeonGreen | 4a,b,c,d | * | 135533 |
| p27 | pCAG_Xph20-mNeonGreen-CCR5TC | pCAG | SV40 | CAG | Xph20 - mNeonGreen-CCR5 ZF-KRAB(A) | Imaging | mNeonGreen | — | * | 135534 |
| p28 | pCAG_Xph15-SNAPf-CCR5TC | pCAG | SV40 | CAG | Xph15 - SNAPf-CCR5 ZF-KRAB(A) | Imaging | Fast kinetics SNAP-tag | 4e,f,g; S6c,d | * | 135536 |
| p29 | pCAG_Xph20-SNAPf-CCR5TC | pCAG | SV40 | CAG | Xph20 - SNAPf-CCR5 ZF-KRAB(A) | Imaging | Fast kinetics SNAP-tag | 6, S8a | * | 135537 |
| p30 | pCAG_Xph20-mEos3.2-CCR5TC | pCAG | SV40 | CAG | Xph20 - mEos3.2-CCR5 ZF-KRAB(A) | Imaging | mEos3.2 | 5; S7 | * | 135532 |
| p31 | pCAG_Xph20-HaloTag-CCR5TC | pCAG | SV40 | CAG | Xph20 - HaloTag-CCR5 ZF-KRAB(A) | Imaging | HaloTag | S8b | * | — |
| p32 | pGP-CMV-GCaMP6f | pCMV | SV40 | CMV | GCaMP6f | Imaging | GCaMP6f | S9, Video 1 | 4 | 40755 |
| p33 | pCMV_Xph15-GCAMP6f | pEGFP-N1 | SV40 | CMV | Xph15 - GCaMP6f | Imaging | GCaMP6f | S9, Video 2 | * | 135538 |
| p34 | pGP-CMV-jGCaMP7f | pCMV | SV40 | CMV | GCaMP7f | Imaging | GCaMP7f | 7 | 5 | 104483 |
| p35 | pCAG_Xph20-GCAMP7f | pCAG | SV40 | CAG | Xph20 - GCaMP7f | Imaging | GCaMP7f | S9 | * | 135539 |
| p36 | pCAG_Xph20-GCAMP7f-CCR5TC | pCAG | SV40 | CAG | Xph20 - GCaMP7f-CCR5 ZF-KRAB(A) | Imaging | GCaMP7f | 7, S9 | * | — |
| p37 | Homer-DsRed | pcDNA3 | SV40 | CMV | Homer - DsRed | Imaging | DsRed | 7, S9 | 6 | — |

**Supplementary Table 3 |** Primers used in this study.

| Name | Seq (5'-3') | Purpose | Used for plasmid |
|---|---|---|---|
| Xop-0612 | gggaattccatatgagctctgtcagttccgtgccg | Xph cloning into pIGc | p06 |
| Xop-0583 | ccgctcgagggtaccggtacggtaattg | Xph cloning into pIGc | p06 |
| Xop-0716 | ggaagatctggcggttcgggcggtagc | Xph cloning into pCAG | p9/10/11/12 |
| Xop-0713 | ccgtgtacattatgcatagtctggcacgtcatac | Xph cloning into pCAG | p9/10/11/12 |
| Xop-0790 | cgcggatccaggcggttcgggcggttcgggaatgtcggtaccgctgac | mIFP cloning into pEGFP-N1 | p13 |
| Xop-0789 | gcgtgtacattatttggactgagactgtgcaaag | mIFP cloning into pEGFP-N1 | p13 |
| Xop-0814 | cccaagcttgccaccatggcggccgaaaaggtcatggagatc | PSD-95-2 cloning into pEGFP-N1 | p13 |
| Xop-0815 | ccgggatcccgggcattgctgggcttggccacc | PSD-95-2 cloning into pEGFP-N1 | p13 |
| Xop-0990 | ggcgctagctccgtgccgaccaaactg | Xph cloning into pCAG-mIRFP670nuc-TEV-xxx | p16/17/18 |
| Xop-1057 | ccgtgtacattaacggtaattgatagaaatc | Xph cloning into pCAG-mIRFP670nuc-TEV-xxx | p16/17/18 |
| Xop-0658 | cggggtaccgcgggcccggggccaccatggcgggcagctctgtcagttccgtg | Xph cloning into pCAG_xxx-eGFP-CCR5TC | p19/20/21 |
| Xop-0607 | ggaagatctactggagccgctaccggtacggtaattgatagaaatc | Xph cloning into pCAG_xxx-eGFP-CCR5TC | p19/20/21 |
| Xop-0788 | cggagatctggtagcggcgtgtctaagggcgaagag | mRuby2 cloning into pCAG_Xph-XXX-CCR5TC | p23 |
| Xop-0787 | ggagctagcagaggagccgctacccttgtacagctcgtccatc | mRuby2 cloning into pCAG_Xph-XXX-CCR5TC | p23 |
| Xop-1054 | gaagatctgtgagcaagggcgaggcag | mScarlet-I cloning into pCAG_Xph-XXX-CCR5TC | p24 |
| Xop-0639 | cggctagccttgtacagctcgtccatgccg | mScarlet-I cloning into pCAG_Xph-XXX-CCR5TC | p24 |
| Xop-1014 | ggcccatgggcagctctgtcagttcc | Xph20-mScarlet-I cloning into pTriex-5 | p25 |
| Xop-1013 | ccgctcgagagagccacctcccacgtgc | Xph20-mScarlet-I cloning into pTriex-5 | p25 |
| Xop-0822 | cggagatctggtagcggcgtgagcaagggcgagg | mNeonGreen cloning into pCAG_Xph-XXX-CCR5TC | p26 |
| Xop-0821 | ggagctagcagaggagccgctacccttg | mNeonGreen cloning into pCAG_Xph-XXX-CCR5TC | p26 |
| Xop-0856 | cggagatctggtagcggcgacaaagactgcgaaatg | SNAPf cloning into pCAG_Xph-XXX-CCR5TC | p28/29 |
| Xop-0855 | ggagctagcagaggagccgctaccgcctgcaggacccagccc | SNAPf cloning into pCAG_Xph-XXX-CCR5TC | p28/29 |
| ODC981 | gctagatctgggagtgcgattaagccagac | mEos3.2 cloning into pCAG_Xph-XXX-CCR5TC | p30 |
| ODC982 | gctgctagctcgtctggcattgtcagg | mEos3.2 cloning into pCAG_Xph-XXX-CCR5TC | p30 |
| ODC983 | ggtgctagcggagctggcgctggagcg | mEos3.2 cloning into pCAG_Xph-XXX-CCR5TC | p30 |
| ODC984 | tctctgtctcgacaagcc | mEos3.2 cloning into pCAG_Xph-XXX-CCR5TC | p30 |
| Xop-1096 | cggagatctggtagcggcgcagaaatcggtactggctttc | HaloTag cloning into pCAG_Xph-XXX-CCR5TC | p31 |
| Xop-1097 | ggagctagcagaggagccgctaccgccggaaatctcgagcgtcg | HaloTag cloning into pCAG_Xph-XXX-CCR5TC | p31 |
| Xop-0726 | cgcggatcccgccaccatggcgggcagctctgtc | Xph cloning into pGP-CMV-GCaMP6f | p33 |
| Xop-0727 | ccgctcgaggctaccgcccgaaccgcc | Xph cloning into pGP-CMV-GCaMP6f | p33 |
| Xop-0658 | cggggtaccgcgggcccggggccaccatggcgggcagctctgtcagttccgtg | Xph cloning into pCAG | p35 |
| Xop-0607 | ggaagatctactggagccgctaccggtacggtaattgatagaaatc | Xph cloning into pCAG | p35 |
| Xop-1129 | cgacgcgtttacttcgctgtcatcatttgtac | GCaMP cloning into pCAG_Xph-XXX-CCR5TC | p35 |
| Xop-1106 | gaagatctgactcatcacgtcgtaagtgg | GCaMP cloning into pCAG_Xph-XXX-CCR5TC | p35/36 |
| Xop-0803 | ccggctagcgctaccgcccgaaccgccgctaccgcccgaaccgcccttcgctgtcatcatttg | GCaMP cloning into pCAG_Xph-XXX-CCR5TC | p36 |

### Supplementary movies

**Supplementary movie 1 and 2**. Spontaneous responses of GCaMP6f (**movie 1**) and Xph15-GCaMP6f (**movie 2**) expressed in hippocampal Banker cultures and recorded at 50 Hz.
